# Chemical perturbations impacting histone acetylation govern colorectal cancer differentiation

**DOI:** 10.1101/2024.12.06.626451

**Authors:** Pornlada Likasitwatanakul, Zhixin Li, Paul Doan, Sandor Spisak, Akhouri Kishore Raghawan, Qi Liu, Priscilla Liow, Sunwoo Lee, David Chen, Pratyusha Bala, Pranshu Sahgal, Daulet Aitymbayev, Jennifer S. Thalappillil, Malvina Papanastasiou, William Hawkins, Steven A. Carr, Haeseong Park, James M. Cleary, Jun Qi, Nilay S. Sethi

## Abstract

Dysregulated epigenetic programs that restrict differentiation, reactivate fetal genes, and confer phenotypic plasticity are critical to colorectal cancer (CRC) development. By screening a small molecule library targeting epigenetic regulators using our dual reporter system, we found that inhibiting histone deacetylase (HDAC) 1/2 promotes CRC differentiation and anti-tumor activity. Comprehensive biochemical, chemical, and genetic experiments revealed that on-target blockade of the HDAC1/2 catalytic domain mediated the differentiated phenotype. Unbiased profiling of histone posttranslational modifications induced by HDAC1/2 inhibition nominated acetylation of specific histone lysine residues as potential regulators of differentiation. Genome-wide assessment of implicated marks indicated that H3K27ac gains at HDAC1/2-bound regions associated with open chromatin and upregulation of differentiation genes upon HDAC1/2 inhibition. Disrupting H3K27ac by degrading acetyltransferase EP300 rescued HDAC1/2 inhibitor-mediated differentiation of a patient-derived CRC model using single cell RNA-sequencing. Genetic screens revealed that DAPK3 contributes to CRC differentiation induced by HDAC1/2 inhibition. These results highlight the importance of specific chemically targetable histone modifications in governing cancer cell states and epigenetic reprogramming as a therapeutic strategy in CRC.

**BRIEF SUMMARY:** HDAC1/2 inhibition promotes colorectal cancer differentiation via gains in H3K27ac, which can be reversed by blocking its acetyltransferase EP300.

## INTRODUCTION

Colorectal cancer (CRC) is the third most common malignancy worldwide and has a rising incidence among the younger population(1). Once diagnosed, metastatic CRC is mainly treated with cytotoxic chemotherapy but responses to these regimens have limited durability. Targeted therapies have shown modest efficacy and inferior results compared to other cancer types harboring identical mutations (e.g., KRAS^G12C^, BRAF^V600E^)(2–10). One explanation for suboptimal targeted therapy responses in CRC is attributed to abnormal cell state plasticity, an inherently complex aspect of cancer development(3,5,11–13). Impaired differentiation is a key mechanism by which cancer cells unlocks phenotypic plasticity(11,14). These features of CRC biology are governed by transcriptional and epigenetic regulation that require further elucidation.

While cancer is conventionally defined by genetic alterations, emerging evidence has highlighted the functional relevance of aberrant epigenetic activity in cancer formation and progression(11,15), including alterations in DNA methylation, histone modifications, nucleosome positioning, and non-coding RNA expression. Epigenetic regulators implicated in cancer biology have been successfully targeted with chemical inhibitors in some settings, offering a promising therapeutic strategy(16,17). In particular, HDAC inhibitors, mainly HDAC1/2 or pan HDAC inhibitors, display potent anti-tumor activity in acute myeloid leukemia (AML) by, in addition to other mechanisms, inducing differentiation(18–20). Five HDAC inhibitors are approved by the FDA for the treatment of hematologic malignancies, either as monotherapy or therapeutic combinations(21). However, HDAC inhibitors have yielded disappointing results when tested in patients with late-stage CRC(22–24).

Inspired by the success of therapeutics that overcome differentiation blocks in leukemias harboring genomic alterations such as BCR-ABL and IDH1/2 (25–27), we aim to identify therapeutic agents that induce CRC differentiation. Recent work from our lab established a critical role for SOX9 in promoting CRC pathogenesis by binding to genome-wide enhancers and reprogramming the epigenetic landscape(14,28). In this setting, SOX9 effectively blocks CRC differentiation by endorsing an aberrant stem cell-like transcriptional program. These findings form the basis of a newly engineered reporter system that emit fluorescent signals from the endogenous *SOX9* and *KRT20* genomic loci of CRC cell lines, broadcasting aberrant stem cell-like and differentiation activity, respectively(29). By applying a chemical screening library to our engineered reporter system, we set out to find and investigate agents that promote CRC differentiation by modulating epigenetic regulation. We aim to increase the likelihood of clinical translation by elucidating the molecular mechanisms by which CRC differentiation is induced, enabling the identification of more selective and potent epigenomic modulators. A stronger molecular evaluation will elucidate the functional connections among specific histone modifications, their impact on gene regulation and chromatin accessibility, and a differentiated cell state phenotype.

## RESULTS

### A chemical library screen identifies the HDAC1/2 inhibitor MRK60 as an inducer of CRC differentiation

Developmental reprogramming and impaired differentiation mediated by SOX9 promotes colorectal cancer (CRC) initiation(14,28). Disrupting SOX9 in mouse models of intestinal neoplasia and human CRC leads to tumor regression by inducing differentiation, which is faithfully indicated by an increase of KRT20 expression similar to levels achieved in normal colon differentiation (30). To effectively produce real-time respective readouts of aberrant stem cell and differentiation activity, we developed a reporter system by genome-editing the endogenous *SOX9* and *KRT20* loci in CRC cell lines to express *mKate2* and *eGFP* fluorescent markers(29) (Figures 1A and S1A). Given the importance of epigenetic regulation in intestinal stem cell biology and dysregulation in CRC, we performed a focused drug screen using 31 well-annotated small molecule inhibitors/degraders of epigenetic regulators applied to two CRC reporter cell lines. The library consists of synthesized compounds that either degrade or disrupt specific enzyme activity of ATP-dependent (e.g., SWI/SNF members) and covalent modifying (e.g., histone deacetylases (HDACs)) chromatin remodelers among other epigenetic regulators (Table S1). The initial screen was performed at a relatively high concentration of 10μM of each compound and resulted in 11 candidates that reduced viability, blocked stem cell activity, and induced differentiation using a permissive cut-off of scoring better than DMSO (Figures 1B and S1B). We next performed a viability secondary screen using the identified 11 compounds at eight different drug concentrations to establish dose-response curves, which yielded six promising candidates (Figure 1C). Five of the top six compounds fell into two classes of epigenetic inhibitors: two compounds targeting SMARCA2/4, subunits of the SWI/SNF complex, and three compounds targeting HDACs.

**Figure 1.**
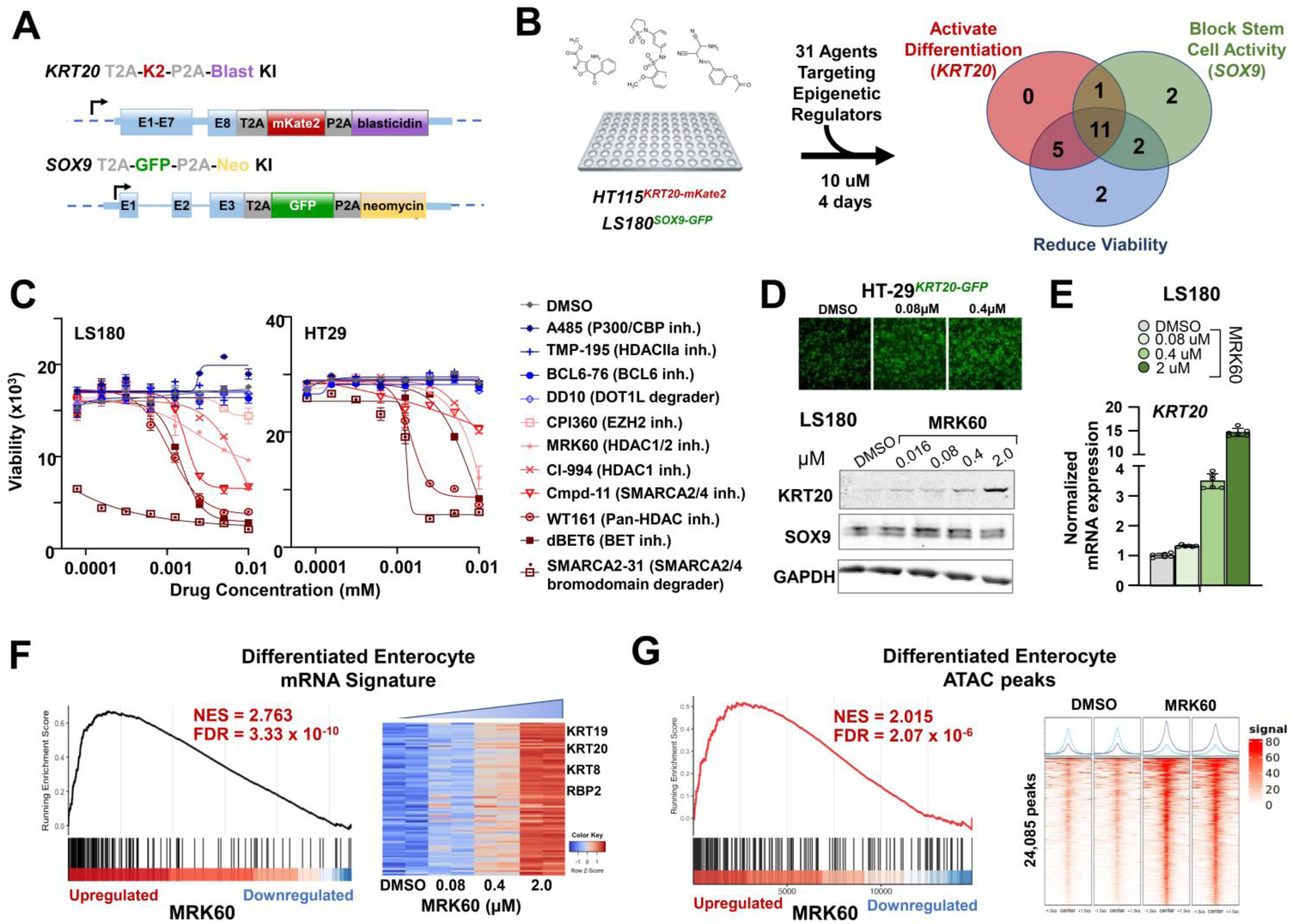
Identification of HDAC1/2 inhibitor MRK60 as a CRC differentiating agent. A. Schematic of endogenous reporter knock-in at *SOX9* and *KRT20* genomic loci. E = exon. B. Schematic of primary drug screen; 3 phenotypic readouts of differentiation (HT115*^KRT20-mKate2^*), stem cell (LS180*^SOX9-GFP^*), and viability across 3 cell lines. Venn diagram shows hits (> DMSO) C. CellTiter-Glo® viability assay showing dose-response curves of LS180 and HT-29 CRC cells treated with 8 doses of 11 drugs at 48 hours D. Representive GFP images of HT-29*^KRT20-GFP^* cells treated with DMSO or indicated concentrations of MRK60 (top); Immunoblot of KRT20, SOX9, and GAPDH in LS180 cells treated with DMSO or indicated concentrations of MRK60 (bottom) E. Normalized mRNA expression of *KRT20* and *SOX9* in MRK60-treated LS180 cells by RT-qPCR. F. Gene set enrichment analysis (GSEA) of differentiated enterocyte signature (Wang, 2020) in RNA-seq profiles of MRK60 (2µM) and DMSO treated HT115 cells; NES = normalized enrichment score and FDR = false discovery rate (left). Heatmap of enriched differentiation genes at indicated MRK60 doses (right). G. GSEA of differentiated enterocyte signature (Wang, 2020) in ATAC-seq profiles of MRK60 (2µM) and DMSO treated HT115 cells; peaks around TSS (±2kb) were annotated and aggregated to the nearest gene (left). Heatmap of open chromatin near differentiation genes (right).

We next evaluated whether the top six compounds could induce CRC differentiation in a dose-dependent fashion by measuring mRNA and protein expression of differentiation marker KRT20 (Figure S1C). Among these six compounds, HDAC 1/2/3 inhibitors, especially the HDAC1/2 selective inhibitor MRK60(31), showed the most consistent and dose-dependent induction of KRT20 at transcriptional and protein levels in CRC cell lines (Figures 1D-E and S1C-G). To confirm the induction of a *bona fide* differentiation program, we performed mRNA and chromatin accessibility profiling of CRC cell lines treated with MRK60 by RNA-seq and assay for transposase-accessible chromatin with sequencing (ATAC-seq), respectively. Gene-set enrichment analyses (GSEA) of transcriptional and chromatin accessibility profiles demonstrated upregulation of a broad intestinal differentiation program(32) (Figures 1F-G and S1G). Consistently, validated intestinal stem cell gene signatures were downregulated upon MRK60 treatment (Figures S1H-I). These data indicate that chemical inhibition of HDAC1/2 by MRK60 promotes differentiation in CRC.

### HDAC1/2 inhibition induces differentiation and impairs tumor growth across CRC models

To confirm that differentiation induced by MRK60 depended on its ability to inhibit HDAC1/2’s enzymatic activity, we synthesized an inactive version of MRK60 (iMRK60) by converting the amine moiety, required for binding zinc in the HDAC enzymatic pocket and inhibiting its catalytic function, to an amide moiety(31) (Figure 2A). Indeed, when applied to CRC cell lines, MRK60 induced differentiation as measured by KRT20 expression, while iMRK60 did not (Figures 2B and S2A-B). In agreement with these results, iMRK60 did not impact CRC proliferation as evaluated by a bioluminescent proliferation assay (Figures 2C and S2C-D).

**Figure 2.**
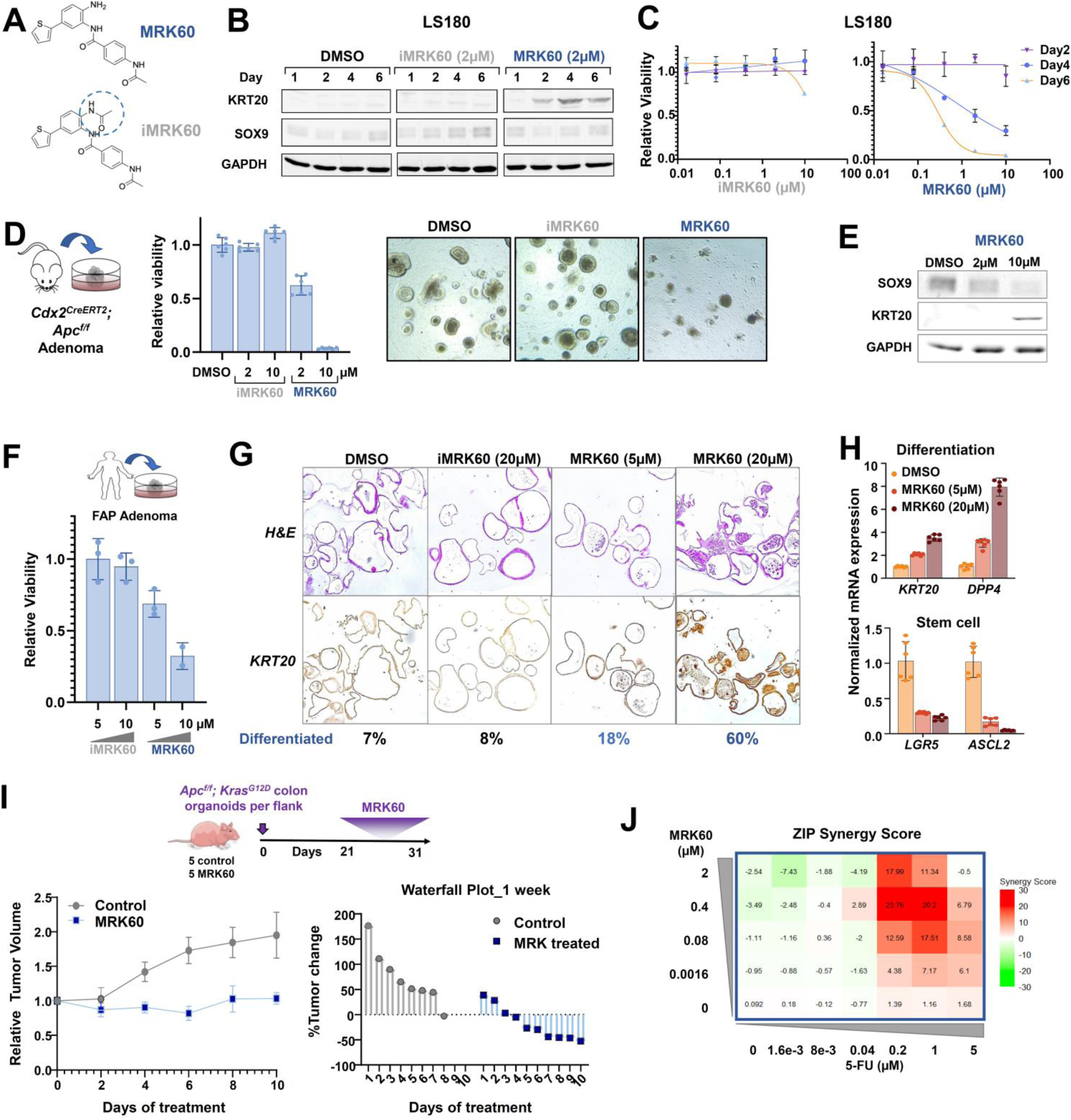
MRK60 induces differentiation and impairs tumor growth across CRC models. A. Structures of MRK60 and its inactive form iMRK60 B. Immunoblots of KRT20, SOX9, and GAPDH (loading control) in LS180 treated with DMSO, iMRK60 (2 µM), or MRK60 (2 µM) at indicated days C. CellTiter-Glo® viability assay of LS180 treated with iMRK60 (left) or MRK60 (right) relative to DMSO at indicated days. D. CellTiter-Glo® viability assay and representative phase contrast images of MRK60 (2 µM), iMRK60 (2 µM), or DMSO treated *Cdx2^CreERT2^; Apc^f/f^* colon adenoma organoids for 4 days. E. Immunoblot of KRT20, SOX9, and GAPDH in *Cdx2^CreERT2^; Apc^f/f^* colon adenoma organoids treated with indicated doses of MRK60. F. CellTiter-Glo® viability assay of patient-derived adenoma organoid from a patient with FAP treated with iMRK60 or MRK60 for 72 hours. G. H&E and KRT20 immunohistochemistry of FAP adenoma treated with iMRK60 or MRK60 for 5 days. Percentage of differentiated organoids (light microscopy) are displayed below images. H. RT-qPCR of differentiation markers (*KRT20*, *DPP4*) and stem cell markers (*LGR5*, *ASCL2*) in FAP adenoma treated with iMRK60 or MRK60 for 72 hours. *ACTB* was used for normalization. I. Schematic of *Apc^f/f^; Kras^G12D^* mouse colon organoid *in vivo* xenograft experiment. Tumor xenograft volume growth curve and waterfall plot of individual tumor responses. J. ZIP synergy score heatmap of HT-29 treated with MRK60 and 5-FU at indicated doses for 5 days.

We also tested the *in vitro* efficacy of MRK60 in three neoplastic colon organoid models. Unlike iMRK60, MRK60 treatment impaired proliferation, induced Krt20 levels, and reduced stem cell gene *Lgr5* expression in premalignant adenoma organoids derived from tdT-labeled Apc^KO^ epithelial cells of *Cdx2^CreERT2^; Apc^f/f^; R26^tdT^* genetically engineered mouse colons following tamoxifen induction (Figures 2D-E and S2F). Similarly, MRK60 treated human adenoma organoids derived from a patient with familial adenomatous polyposis (FAP) displayed reduced viability which, together with increased KRT20 levels and reduced LGR5 expression, demonstrated a differentiation phenotype not observed with iMRK60 treatment (Figure 2F-H). Less pronounced than the 75% reduction in viability observed in FAP adenomas, MRK60 reduced viability of patient-derived CRC organoid by 50% at the highest dose (Figure S2G).

To evaluate the *in vivo* anti-tumor properties of MRK60, we treated nude mice harboring flank tumors derived from transplanted mouse Apc^KO^; Kras^G12D^ colon organoids. Mice were randomized to vehicle control or daily MRK60 treatment by intraperitoneal injection 21 days post-implantation. All MRK60 treated mice had either tumor shrinkage or stability after 10 days whereas control mice displayed tumor doubling during the same time frame (Figure 2I). Using the *Lgr5^Cre^; Apc^F/F^;R26^tdT^* genetically engineered mouse model of intestinal polyposis, we found that *in vivo* treatment of MRK60 induced *Krt20 levels* that corresponded to a modest increase in protein expression by immunohistochemistry (IHC) (Figure S2H).

We next examined if existing CRC therapeutics may combine with MRK60 to improve differentiation and anti-tumor activity. In combination with MRK60, we treated two CRC cell lines with three different chemotherapeutics (5-fluorouracil (5-FU), oxaliplatin, and the active metabolite of irinotecan SN38) and two targeted agents (Regorafenib, Trifluridine/Tipiracil (TAS102)), and found that only 5-FU synergized with MRK60 (Figure 2J and Figure S2I). The ZIP score, a statistical algorithm that calculates drug syndergy(33), indicated synergy was achieved when 5-FU reached a concentration of 1µM (Figure 2J), consistent with the concentrations of 5-FU and MRK60 combinations that maximally induced KRT20 expression as assessed by the endogenous differentiation reporter system (Figure S2J). Treatment of our patient-derived CRC organoid showed an additive anti-viability effect of 5-FU and MRK60, which corresponded to elevated expression of KRT20 and suppression of stem cell marker *LGR5* (Figure S2K-M).

### MRK60 induces differentiation via on-target HDAC1/2 inhibition

We next sought to investigate whether the differentiation effect of MRK60 is mediated by on-target HDAC1/2 inhibition via biochemical, genetic, and chemical inhibitor assays (Figure 3A). To validate that MRK60’s effect is through physical engagement of HDAC1 and HDAC2, we evaluated protein interactions of MRK60 using an unbiased proteomic assessment. We generated biotinylated MRK60 and iMRK60 (see Methods), separately incubated them with CRC protein lysates, then performed streptavidin-bead pulldowns followed by mass spectrometry (Figure 3B). This experiment showed that MRK60 displays a strong and preferential physical interaction with HDAC1 and HDAC2 relative to iMRK60 in two independent CRC (Figures 3B-C and S3A-B). Indeed, biochemical pull-down assays confirmed a specific interaction between active MRK60 and HDAC1 (Figure 3D). In addition to HDAC1/2, eIF-2-alpha kinase activator GCN1 (GCN1) was found to interact with MRK60 in both cell lines; however, GCN1 knockout by CRISPR/Cas9 did not induce differentiation, suggesting it is likely not involved in MRK60-induced differentiation (Figures 3B and S3B-C). Individual knockout of HDAC1 or HDAC2 using CRISPR/Cas9 led to a modest induction in differentiation in CRC cell lines as measured by KRT20 levels (Figure S3D); however, the effect wasn’t as strong as MRK60, which simultaneously inhibits HDAC1 and HDAC2 catalytic function.

**Figure 3.**
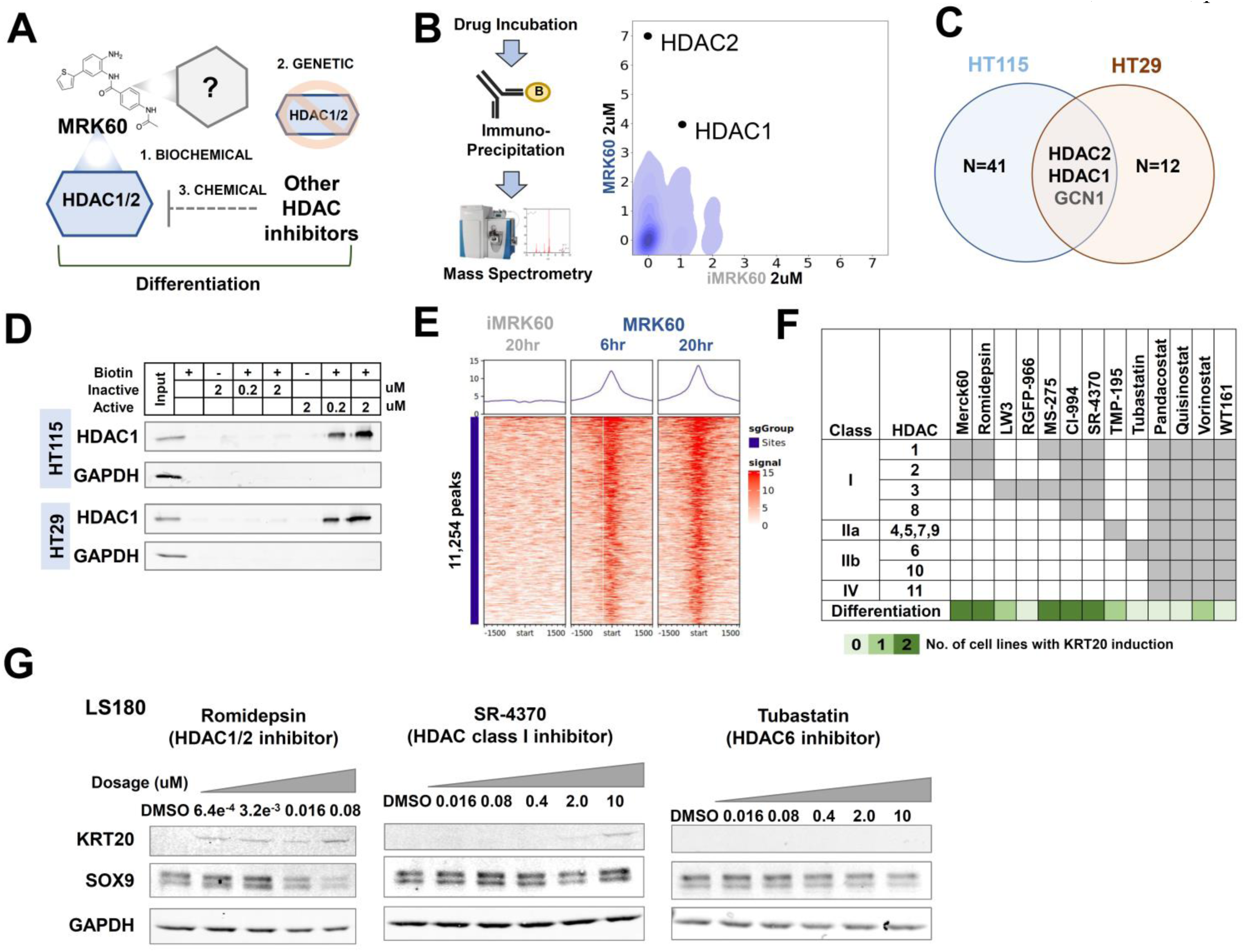
MRK60-induced differentiation is mediated by on-target HDAC1/2 inhibition. A. Schematic of strategies to validate on-target effect of MRK60 on HDAC1/2 inhibition B. Schematic of MRK60 IP followed by mass spectometry workflow (left); density plots of unique peptides bound to biotinylated MRK60 (y-axis) or iMRK60 (x-axis) in HT-29 cells (right). C. Venn diagram showing proteins detected by co-IP mass spectometry in HT-115 and HT-29. D. Immunoblots of HDAC1 and GAPDH in HT-115 and HT-29 following co-IP with biotin control, iMRK60, biotinylated iMRK60, MRK60, or biotinylated MRK60. E. Heatmaps of iMRK60 and MRK60 binding at genomic loci co-bound by HDAC1/2 using Chem-seq protocol in live CRC cells F. A table displaying ability of 13 inhibitors with indicated HDAC inhibitory profiles and specificity to induce differentiation in CRC cell lines upon treatment G. Immunoblots of KRT20, SOX9, and GAPDH in LS180 treated with indicated HDAC inhibitors

To characterize MRK60-HDAC1/2 complex DNA engagement, we developed and deployed a chemical probe-based sequencing method called Chem-seq(34), which identifies regions of the genome that correspond to chemical probe interactions with DNA-binding proteins. Applying this approach, CRC cells were separately treated with biotinylated MRK60 and iMRK60, followed by streptavidin pulldown and DNA sequencing. Biotin-MRK60, but not biotin-iMRK60, associated with HDAC1/2 peaks throughout the genome, with greater binding intensity over time (6h and 20h) (Figures 3E and S3E).

Finally, we used a chemical approach to interrogate whether inhibition of specific HDACs is associated with the differentiation phenotype in CRC. We treated two CRC cell lines with a series of HDAC inhibitors including six class I HDAC inhibitors (HDAC1,2,3,8), two class II HDAC inhibitors (HDAC 4-7,9,10), and four pan-HDAC inhibitors (Figure 3F). The differentiation phenotype, denoted by KRT20 induction detected by immunoblots, correlated most with treatment of class I HDAC inhibitors, particularly those annotated to inhibit HDAC 1, 2, or 3 (Figures 3F-G and S3F). MRK60, Romidepsin(35), MS-275(36), Cl-994(37), and SR-4370(38) were the best performing HDAC inhibitors, whereas HDAC3 inhibitors LW3(39) and RGFP-966(40) did not consistently induce KRT20 (Figures 3F-G and S3F). Class II and pan-HDAC inhibitors similarly did not reliably induce KRT20 (Figures 3F-G and S3F). HDAC deacetylases can operate individually or as part of co-repressor complexes, such as CoREST. Treatment of CRC cells with UM-171(41), a CoREST degrader, did not induce differentiation, suggesting that co-repressor complexes may not play an important role in the differentiation process (Figure S3G).

### Integrative analyses of HDAC1/2 inhibition on chromatin accessibility and gene regulation

To define the mechanism(s) by which MRK60 promotes CRC differentiation, we analyzed gene expression profiles from CRC cells treated with MRK60 and identified 6,606 differentially expressed genes (DEGs) with an almost equal split between up- and downregulated genes, in alignment with prior work(42). Consistent with our earlier results (Figure 1F), differentiation markers were among the top upregulated DEGs (e.g., KRT8, KRT19, and KRT20); in contrast, stem cell genes (e.g., *ASCL2*) were enriched among the 3227 downregulated genes (Figure 4A). Gene ontology demonstrated that MRK60 induces gene programs associated with developmental maturation and suppressed genes associated with growth regulators as suggested by disenabling DNA replication, methylation, and RNA splicing processes being significantly downregulated (Figure S4A).

**Figure 4.**
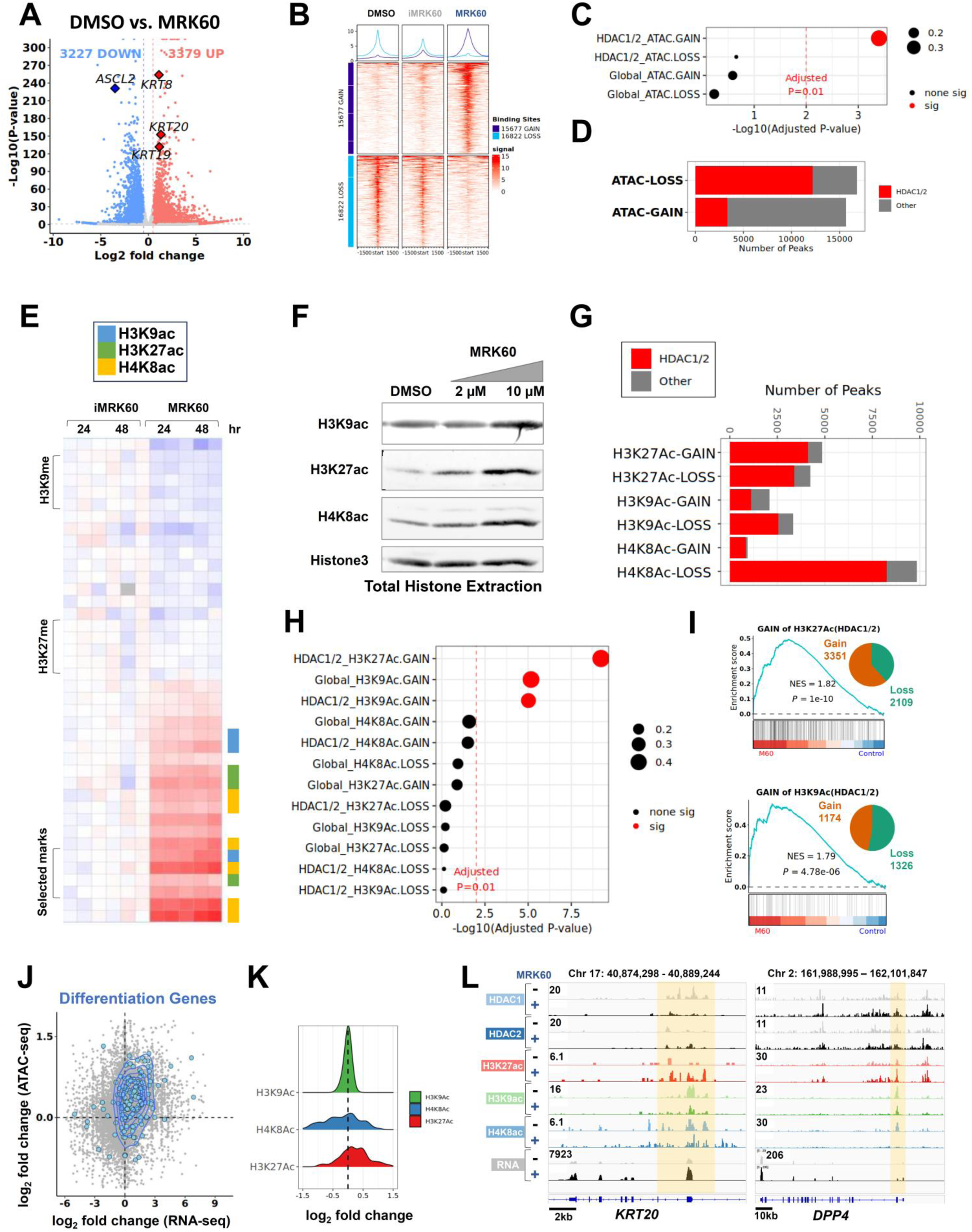
Gains in H3K27ac mediated by HDAC1/2 inhibition are associated with CRC differentiation. A. Volcano plot of DEGs in HT-115 treated with DMSO and MRK60 by RNA-seq. Genes with adjusted *P*-value < 0.05 are highlighted; downregulated (blue) and upregulated (red) in MRK60 B. Heatmaps of ATAC-seq peaks in HT-115 cells treated with DMSO, iMRK60, and MRK60; differential accessibility peaks with adjusted *P*-value < 0.05 and absoluate log_2_ fold change >0.5. C. GSEA dot plot of MRK60 DEG by RNA-seq associated with global and HDAC1/2-bound ATAC peak gains and losses. Dot size indicates the proportion of enriched genes in entire gene set. Adjusted *P*-value = 0.01 cutoff is indicated by dashed red line. D. Bar plots of MRK60-induced gained and lost ATAC peaks; HDAC1/2 bound subset marked in red E. Heatmap of gained (red) and lost (blue) histone post-translational modifications (PTMs) in HT-29 cells by mass spectometry; H3K9ac (blue), H3K27ac (green), and H4K8ac (yellow). F. Histone-enriched immunoblots of H3K9ac, H3K27ac, H4K8ac, and H3 in MRK60-treated HT-115 G. Bar plots MRK60-induced of gained and lost peaks of H3K27ac, H3K9ac and H4K8ac; the proportion bound by HDAC1/2 indicated in red. H. GSEA dot plot of MRK60 DEG by RNA-seq associated with global and HDAC1/2-bound changes in H3K27ac, H3K9ac and H4K8ac. I. GSEA of MRK60-induced DEG in HT-115 by RNA-seq among MRK60-induced gained H3K27ac (top) and H3K9ac (top) bound by HDAC1/2; NES, normalized enrichment score. J. Scatter plot of MRK60-induced log_2_-fold changes in chromatin accessibility by ATAC-seq (y-axis) and gene expression by RNA-seq (x-axis); differentiation genes and countour map in blue K. Density plot of MRK60-induced distribution of H3K27ac, H3K9ac and H4K8ac among MRK60 upregulated differentiation genes. L. IGV tracks for HDAC1, HDAC2, H3K27ac, H3K9ac, H4K8ac, and RNA-seq at genomic loci of differentiation genes *KRT20* and *DPP4* in HT-29 cells +/- MRK60.

Histone acetylation regulates, among other processes, DNA accessibility. We therefore evaluated how HDAC1/2 inhibition impacts chromatin access by ATAC-seq, aiming to relate these results to the observed gene expression changes. Similar to changes in mRNA profiles, there were an equal number of gained and lost chromatin accessibility regions upon MRK60 treatment, with 15,677 newly accessible compared to 16,822 closed regions (Figure 4B). While global gained chromatin regions related to upregulated genes more than lost regions associated with downregulated genes, neither reached statistical significance (Figure 4C), which may be due to the larger magnitude of changes in accessibility compared to gene expression as well as indirect effects of HDAC1/2 inhibition. To ask if HDAC1/2- bound regions better resolve the direct effect of MRK60 with respect to chromatin accessibility and gene expression changes, we performed HDAC1/2 chromatin-immunoprecipitation followed by DNA sequencing (ChIP-seq). 72.5% of newly closed chromatin overlapped with HDAC1/2 binding sites compared to the 21.1% of gained ATAC peaks (Figure 4D), which was unexpected as blocking deacetylation is typically associated with opening chromatin. Nevertheless, the 3,308 gained ATAC peaks bound by HDAC1/2 significantly associated with transcriptional upregulation by GSEA, whereas the 12,204 lost ATAC peaks bound by HDAC1/2 did not correlate with downregulated genes (Figures 4C and S4B-C). Also, MRK60-bound regions by Chem-seq strongly overlapped with HDAC1/2-bound regions by ChIP-seq, indicating that MRK60’s ability to impact gene regulation broadly and induce differentiation specifically through direct engagement of HDAC1/2 without disrupting its interaction with chromatin (Figure S3E). Taken together, these data signify that direct gene regulatory effects of HDAC1/2 inhibition operate through opening chromatin at HDAC1/2-bound regions to upregulate transcription.

### Chemical inhibition of HDAC1/2 leads to gains in acetylation of specific histones

HDACs impact gene regulation by removing acetyl groups from different lysine residues of histone and non-histone proteins. To connect the gene expression and chromatin accessibility changes of MRK60 to HDAC1/2’s direct function and identify the specific lysine residues associated with differentiation induction, we performed H2, H3 and H4 histone posttranslational modification (PTM) profiling by mass spectrometry(43). After histone specific immunoprecipitation, we interrogated 79 different PTMs including (methylation, acetylation (ac), and ubiquitinoylation) in two CRC lines treated with MRK60 and iMRK60 (Figure 4E). As expected, many H3 and H4 lysine residues displayed greater acetylation upon MRK60 treatment, with H3K27ac, H3K9ac, and H4K8ac showing the greatest increases, nominating them as potential histone marks to further interrogate (Figure 4E). To validate these findings, we measured protein levels of these acetylated histones in CRC cell lines treated with MRK60 using histone-enrichment immunoblots, observing a dose-dependent increase in their levels (Figures 4F and S4D).

### Acetylation of H3K27 mediated by HDAC1/2 inhibition is associated with CRC differentiation

To investigate locus-specific histone acetylation and correlate these with changes in chromatin accessibility and gene expression upon HDAC1/2 inhibition, we profiled H4K8ac, H3K27ac and H3K9ac in CRC cell lines treated with MRK60 by CUT&RUN(44). Quality control by fingerprint analysis(45) showed all antibody-treatments enriched sufficiently, peak signal separated from background (Figure S4E), and replicates were consistent (Figure S4G). The genomic regions with MRK60-induced changes in histone acetylation largely overlapped with HDAC1/2 binding, with the following distributions: active gene body (48.3%), distal intergenic (30.8%) and promoters (28.0%) (Figure S4F). Indeed, changes in H3K27ac, H3K9ac, and H4K8ac were predominantly found at HDAC1/2-bound regions (Figure 4G), suggesting a direct relationship between HDAC1/2 and these chromatin-associated histone PTMs. This pattern was also observed with CRC cells treated with Romidepsin, a second HDAC1/2 inhibitor (Figure S4H). A comparable number of genomic regions gained and lost H3K27ac and H3K9ac. Surprisingly, H4K8ac was predominantly lost upon HDAC1/2 inhibition with either drug (Figures 4G and S4H-J), which was unexpected given the greater abundance of H4K8ac (Figure 4E-F). We confirmed that MRK60 did not induce changes in cellular localization of these acetylated histones by immunofluorescence (Figure S4I).

We next asked how changes in histone acetylation related to chromatin accessibility. The HDAC1/2- bound genomic regions that opened their chromatin upon HDAC1/2 inhibition corresponded to gained H3K27ac and H3K9ac marks, which was not the case for areas that lost chromatin accessibility upon HDAC1/2 inhibition. H4K8ac was significantly reduced at genomic regions that gained and lost chromatin access, suggesting no relationship with DNA accessibility (Figure S4K).

Finally, we integrated histone acetylation, chromatin accessibility and transcriptional profiles to generate a comprehensive molecular portrait. Upon HDAC1/2 inhibition, we found that HDAC1/2-bound regions that gained H3K27ac and H3K9ac were significantly associated with transcriptional upregulation (Figure 4H-I). Notably, open chromatin and upregulated genes associated with differentiation were most significantly linked to HDAC1/2-bound regions that gained H3K27ac but not H3K9ac or H4K8ac (Figure 4J-L). This suggests that inhibition of HDAC1/2 deacetylase activities could lead to direct accumulation of H3K27ac near differentiation genes, facilitating chromatin access and gene expression. In contrast, closed chromatin and downregulated genes associated with stem cell activity were most significantly connected to HDAC1/2-bound regions that lost H3K27ac and, to a lesser extent, H3K9ac and H4K8ac (Figure S4L-M). Together, these findings support a model whereby chemical inhibition of HDAC1/2 leads to accumulation of H3K27ac at differentiation genes in CRC.

### Histone acetyltransferase EP300 is required for CRC differentiation mediated by HDAC1/2 inhibition

To investigate if the accumulation of H3K27ac following HDAC1/2 inhibition is required for CRC differentiation, we disrupted EP300, a principal acetyltransferase responsible for depositing acetyl groups on H3K27, asking if the differentiation phenotype could be reversed. We started by exposing MRK60 treated CRC cell lines to JQAD1, a recently developed EP300 degrader by our group(46). We found that JQAD1 reversed MRK60-induced KRT20 levels in a dose-dependent fashion by immunoblot (Figure 5A). This corresponded to reduced H3K27ac levels in CRC cells treated with JQAD1 and MRK60 by H3-enriched immunoblot (Figure 5B). Using our endogenous differentiation reporter system (HT29*^KRT20-GFP^*), we observed that the approximately 3-fold increase in the percentage of KRT20^+^ cells induced by MRK60 was reduced 2-fold upon concomitant JQAD1 treatment, almost returning to baseline levels (Figure 5C). By analyzing the results of our recently published genetic screen targeting epigenetic regulators(29), we found that EP300 knockout was one of the strongest perturbations supporting a stem cell-like state as measured by our dual reporter system (Figure 5D).

**Figure 5:**
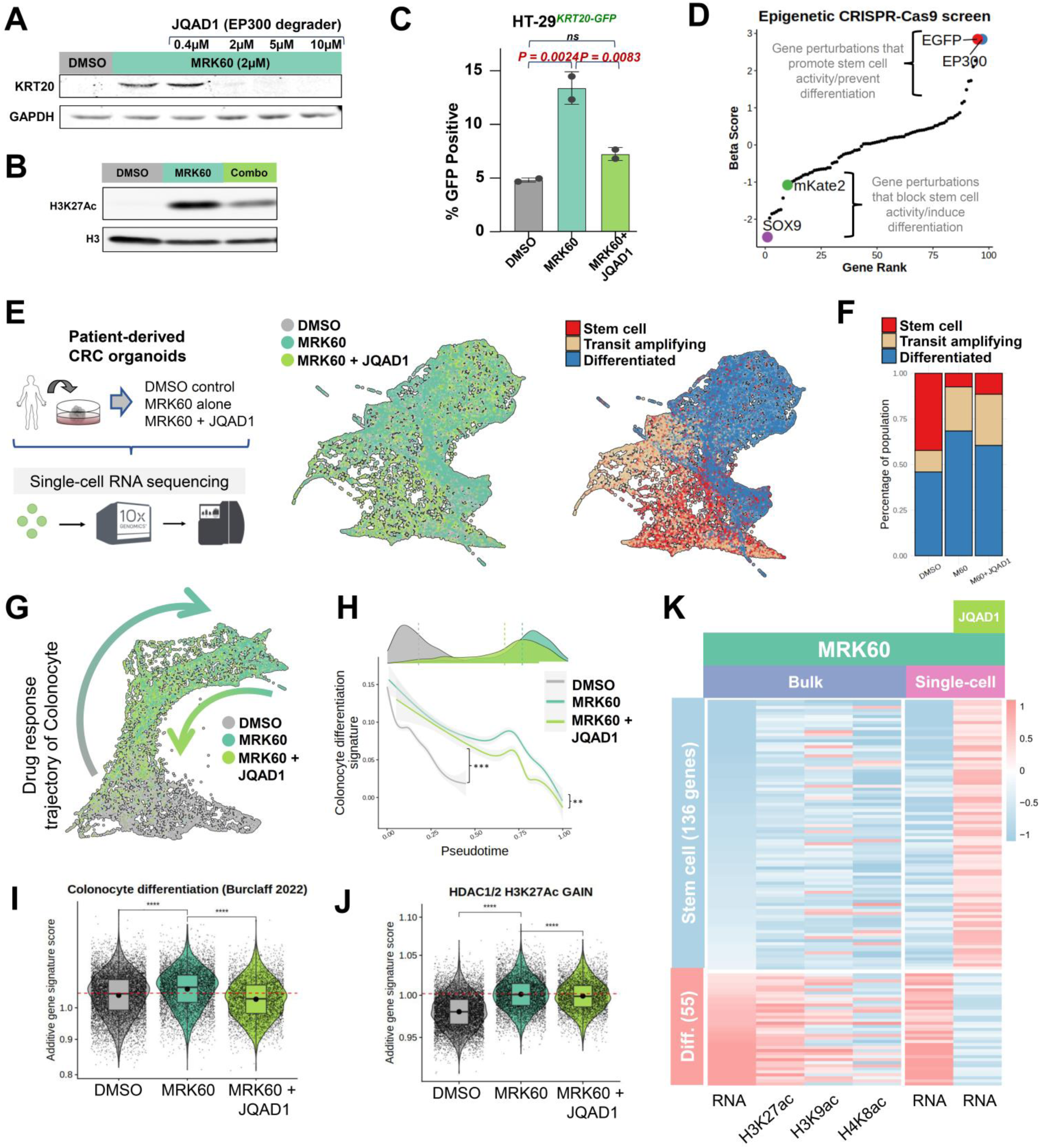
H3K27ac is required for differentiation reprogramming by HDAC1/2 inhibition. A. Immunoblot of KRT20 and GAPDH proteins in LS180 treated with MRK60 (2 µM) and indicated concentrations of JQAD1 (0.4 - 10 µM). B. Histone-enriched immunoblot of H3K27ac and H3 in HT-29 treated with MRK60 (5 µM) +/- JQAD1 (10 µM). C. Quantification of GFP+ percentage in HT29*^KRT20-GFP^* reporter cell line treated with MRK60 (5 µM) +/- JQAD1 (10 µM). D. Ranked log_2_ fold change of sgRNA distribution in GFP^high^ (perturbation promotes differentiation) and mKate2^high^ (perturbation promotes stem cell) sorted cell fractions of published CRISPR-Cas9 screen targeting epigenetic regulators (78 genes 542 sgRNAs) using HT29*^SOX9-mKate2/KRT20-GFP^* reporter cell line^16^; select perturbations indicated in colored circles. E. Schematic of scRNA-seq experiment in patient-derived CRC organoids treated with DMSO, MRK60, or MRK60+JQAD1 combination (left). UMAPs of scRNA-seq colored by treatment group (middle) or cell type (right). F. Proportion of cell types in CRC organoids treated with DMSO, MRK60, or MRK60+JQAD1. G. Trajectory analysis of colonocytes in scRNA-seq experiment colored by treatment group. H. Expression of colonocyte differentiation signatures along drug response trajectory. Density plot shows histogram of cells in different treatment along pseudotime. ** *P* = 0.0042; *** *P* = 0.0002 I. Violin plot of colonocyte differentiation gene signature in scRNA-seq experiment by treatment groups. **** *P* = 7.0 x 10^-6^ (left), **** *P* = 8.4 x 10^-8^(right) J. Violin plot of gene signatures associated with MRK60-induced gains in H3K27ac bound by HDAC1/2 in scRNA-seq experiment by treatment groups. **** *P* = 5.8 x 10^-10^ (left), **** *P* = 2.1 x 10^-5^(right) K. Integrative heatmap of expression changes in stem cell and differentiation genes associated with H3K27ac, H3K9ac, and H4K8ac in bulk and single-cell RNA data sets with MRK60 and/or JQAD1.

Cell type heterogeneity is not easily resolved by bulk RNA analysis. To address this limitation and decipher how stem cell and differentiation cell states are regulated by histone acetylation in CRC, we performed single-cell RNA sequencing (scRNA-seq) on patient-derived CRC organoids treated with DMSO control, MRK60 and MRK60+JQAD1 (Figure 5E). We obtained 33,331 high-quality single cells that were classified into 3 major epithelial cell types, including intestinal stem cell (ISC, 24.9%); transit amplifying (TA, 19.1%); absorptive colonocyte (ACC, 55.9%) (Figure 5E). To corroborate our clustering results, we compared our datasets against a well-annotated human reference scRNA-seq dataset(47) by using the Harmony integration tool(48), and confirmed similar clustering patterns. All clusters contained cells from each treatment group, suggesting the absence of dissociation bias, and each cell type uniquely expressed expected cell-defining marker genes (Figure S5A), supporting strong quality metrics. Overall, we detected an average of 3,256 unique genes and 10,142 transcripts per cell (Figure S5B).

MRK60 treatment reduced the proportion of ISC and TA cells and correspondingly increased the percentage of differentiated colonocytes (Figure 5F-H). The proportion of cells entering G1 cell cycle arrest also increased (Figure S5C), consistent with differentiated cells entering a post-mitotic state. MRK60 treatment upregulated differentiation genes that more strongly associated with HDAC1/2-bound H3K27ac gains than those of H3K9Ac and H4K8ac (Figures 5I-K and S5F-G), consistent with our previous results (Figures 4H-L).

To evaluate if H3K27ac is responsible for colonocyte differentiation, we degraded EP300 using JQAD1 and asked if the differentiation phenotype induced by MRK60 is reversed. The scRNA-seq results showed that the combination treatment could partially reverse the effect of MRK60 treatment, increasing the proportion of ISC and TA cells while reducing the percentage of differentiated colonocytes (Figure 5F). Compared to DMSO, we identified 3,006 and 3,831 unique DEGs in MRK60 and combination treatment, respectively (false discovery rate [FDR] < 0.01, log2 fold change > 1.5) (File 1). DEGs were partially reversed by combination treatment compared to MRK60 treatment alone (Figure S5D). We found that the DEGs identified by scRNA-seq were highly correlated with those identified by bulk RNA-seq (Figure S5E-G), reflecting robust results across different sequencing technologies and CRC models.

We constructed a drug response trajectory for colonocytes and observed that cells treated with MRK60 treatment projected along a differentiation continuum, whereas the addition of JQAD1 pushed cells back towards stem and transit amplifying cell states (Figure 5G-H). JQAD1 treatment downregulated colonocyte differentiation gene expression signatures promoted by HDAC1/2 inhibition (Figure 5H&I). Furthermore, JQAD1-mediated downregulated genes most strongly associated with HDAC1/2-bound regions of H3K27ac and, to a lesser extent, H3K9ac (Figures 5J-K and S5G). In line with these findings, MRK60 and JQAD1+MRK60 treatments showed opposite effects on stem(49) and differentiation(47) genes signatures by GSEA (Figures 5I and S5H). Collectively, these results suggest that chemically induced gains in H3K27ac and CRC differentiation can be reversed by degrading its writer EP300.

### DAPK3 is induced by HDAC1/2 inhibition and a functional regulator of differentiation

In addition to direct activation of differentiation genes by gains in H3K27ac, we wondered if HDAC1/2 inhibition induced genes that facilitated differentiation. To narrow the list of potential targets to interrogate, we identified genes that upon treatment of CRC cells with MRK60 were (1) bound by HDAC1/2 (both HDAC1/2 Cut&Run and MRK60 Chem-seq), (2) associated with gains in H3K27ac, and (3) displayed upregulated gene expression (Figure 6A). We then performed a pooled CRISPR-Cas9 screen and asked which of the perturbed genes reversed MRK60-induced loss of viability and differentiation using HT-29*^KRT20-GFP^* reporter line (Figures 6B-C). Among 27 short-listed factors, DAPK3 (Death Associated Protein Kinase 3), a chromatin-(50) and apoptosis-associated kinase(51,52) that functions in cell cycle regulation and cell survival(52–55), emerged as a potential regulator of CRC viability and differentiation. *DAPK3* targeting sgRNAs ranked 3^rd^ and 4^th^ in disrupting MRK60 mediated loss of viability and differentiation induction, respectively (Figure 6C and S6A).

**Figure 6:**
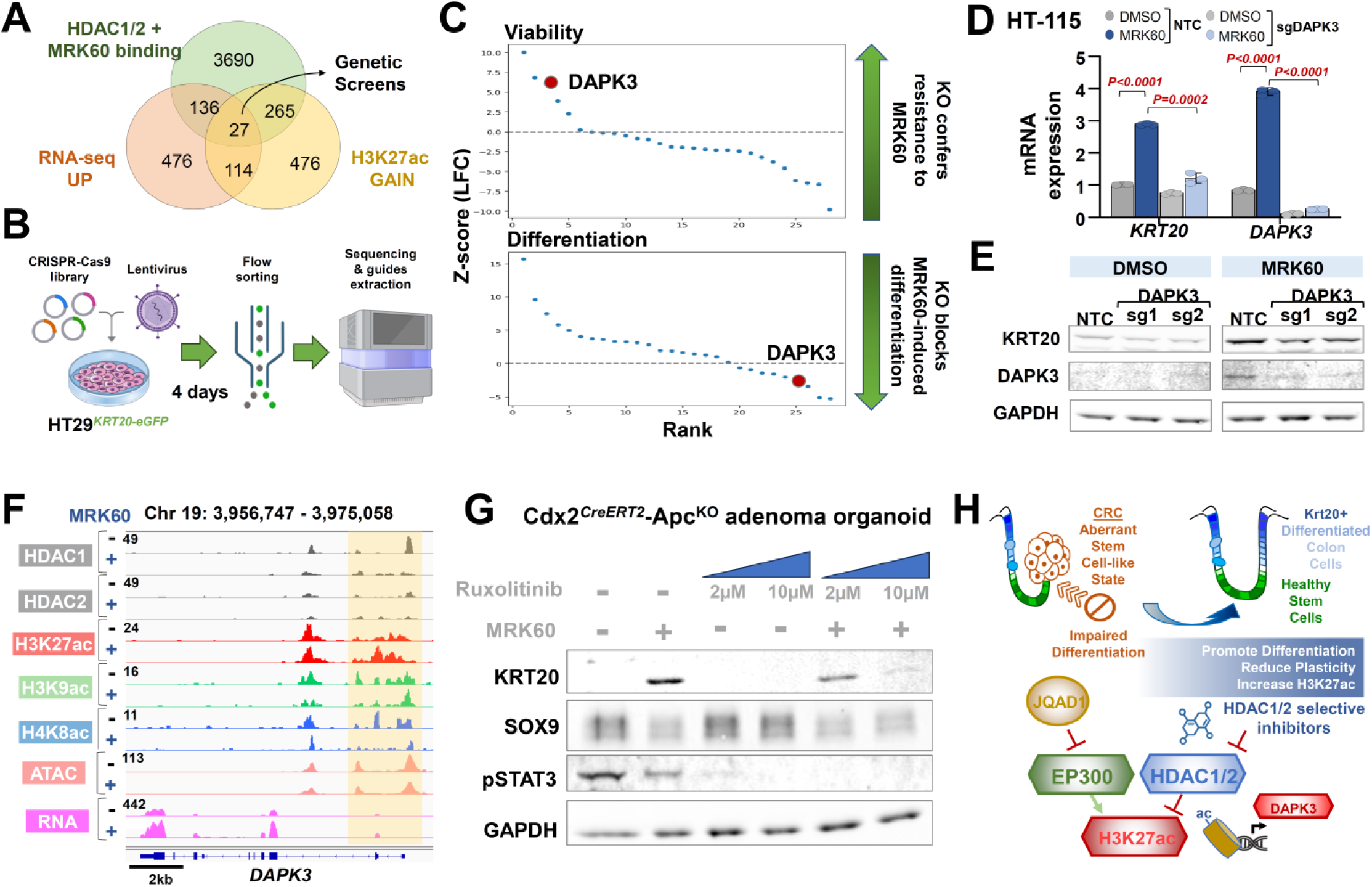
DAPK3 is upregulated and required for differentiation mediated by HDAC1/2 inhibition. A. Venn diagram showing genes bound by HDAC1/2 and MRK60, upregulated by MRK60, and gained H3K27ac upon MRK60 treatment. B. Schematic diagram showing CRISPR-Cas9 genetic screen in HT29*^KRT20-GFP^* reporter cells. C. Ranked normalized Z-scored log_2_ fold change (LFC) plots of sgRNA targeting 27 genes in HT29*^KRT20-GFP^* reporter screen with respect to viability (top) and GFP/KRT20 (bottom) outputs D. mRNA and (E) protein expression of KRT20 and DAPK3 in HT115 cells engineered to suppress DAPK3 by CRISPRi knockdown treated with 2µM MRK60 for 72 hours. F. Integrative Genomic Viewer (IGV) snapshot depicting HDAC1/2 binding, H3K27ac, H3K9ac, and H4K8ac CUT&RUN peaks and ATAC-seq profiles at DAPK3 genomic locus. G. Immunoblots of *Cdx2^CreERT2^; Apc^f/f^; R26^tdT^* adenoma mouse organoids treated with 5µM MRK60 and indicated doses of Ruxolitinib at 72 hours. H. Schematic diagram showing a proposed differentiation mechanism mediated by HDAC1/2 and H3K27ac

HDAC1/2 inhibition transcriptionally activated DAPK3 in each CRC cell line and organoid model tested (Figures 6D-E and S6B-E). At the *DAPK3* genomic locus, HDAC1/2 inhibition increased H3K27ac and RNA levels, reduced HDAC1/2 binding and H4K8ac, and unaffected H3K9ac and chromatin accessibility by ATAC-seq (Figure 6F). DAPK3 suppression by CRISPRi partially reversed differentiation induced by HDAC1/2 inhibition in two CRC cell lines (Figures 6D-E and S6B-E), validating our screening results. In agreement with these findings, CRC cell lines with lower DAPK3 expression were more resistant to MRK60 in BROAD’s CTD^2 drug screen (Figure S6F, *P = 0.045*).

JAK1/2 kinase inhibitor Ruxolitinib is annotated to inhibit DAPK3 at Kd of 89nM (Kd for JAK1, JAK2, and JAK3 are 3.4nM, 0.036nM, and 2nM, respectively)(56,57). DAPK3 and JAK3 are the only two Ruxolitinib targets upregulated by MRK60 (Figure S6G). We therefore tested if chemical inhibition of DAPK3 by Ruxolitinib could reverse the CRC differentiation phenotype induced by HDAC1/2 inhibition. Dose-dependent Ruxolitinib treatment reduced differentiation induction by HDAC1/2 inhibition in Cdx2-Apc^KO^ adenoma organoids and CRC cell lines as measured by KRT20 levels (Figures 6G and S6D-E); consistently, stem cell markers displayed elevated expression upon combination treatment (Figure S6D-E). JAK/STAT pathway genes did not show mRNA expression patterns associated with differentiation in Cdx2-Apc^KO^ adenoma organoids or HT-115 CRC cells treated with MRK60 and/or Ruxolitinib (Figures S6H-I); in fact, JAK3 and other pathway genes were downregulated upon HDAC1/2 inhibition. Furthermore, genetic knockout of JAK1, JAK2, JAK3, and STAT3 did not impact KRT20 induction by MRK60 (Figures S6J-K), suggesting that Ruxolitinib’s effect on blocking MRK60-induced CRC differentiation is mediated through DAPK3 inhibition. Consistent with its effect on differentiation, the combination of MRK60 and Ruxolinitib displayed dose-dependent anti-synergism with respect to viability in a CRC cell line by ZIP synergy score calculations (Figure S6L). Collectively, these data indicate that DAPK3 is a functional component of HDAC1/2 inhibition mediated CRC differentiation (Figure 6H).

## DISCUSSION

Genetic mutations and epigenetic alterations promote CRC initiation, progression, and therapy resistance. These alterations promote a stem cell-like state with aberrant gene expression programs that confer CRC plasticity(14,58–61). While the sequence of genetic alterations involving APC, p53, SMAD4, and KRAS are well established (62), the functional characterization of epigenetic dysregulation in CRC plasticity is incomplete and continues to grow in importance(63,64). Garnering attention as a recently appreciated hallmark of cancer, nongenetic and epigenetic mechanisms have been described to underlie CRC initiation and therapy resistance(11). We established a dual endogenous reporter system by genome engineering *SOX9* and *KRT20* loci in CRC cell lines to broadcast fluorescent readouts of aberrant stem cell (i.e., pro-cancer) and differentiation (i.e., anti-cancer) activity, respectively(29). With this system, we strived to capture functional readouts of CRC plasticity that are governed by epigenetic regulation. By screening inhibitors or degraders of epigenetic regulators using this platform, we set out to identify agents that reduce aberrant stem cell activity and induce CRC differentiation, borrowing from a successful therapeutic paradigm of leukemia (26).

We uncovered HDAC1/2 inhibitors and SMARCA2/4 degraders as promising candidates using our reporter system and viability screens and focused on HDAC1/2 inhibitors based on their consistency in activating differentiation across models and clinical applicability for treating patients with hematological malignancies. There are five FDA-approved HDAC inhibitors for the treatment of T-cell lymphoma and multiple myeloma. Three pan-HDAC inhibitors (Vorinostat, Belinostat, Panobinostat), one class I HDAC inhibitor (Romidepsin), and one class I/II HDAC inhibitor (Tucidinostat). Vorinostat and Romidepsin have also been evaluated in phase II clinical trials as monotherapy in several solid cancers, including advanced CRC patients(24,65). One factor limiting their clinical success is the risk of cardiac and hematologic side effects likely due to protein post-translational modifications mediated by HDAC1/2 inhibition. By identifying the subset of changes in protein acetylation that promote differentiation and engender anti-tumor activity, we are able to develop new strategies to treat CRC while avoiding these untoward side effects. We narrowed our search for HDAC1/2 inhibitor-mediated differentiation changes to histones based on at least two concepts: (1) a differentiation phenotype is likely governed by transcriptional reprogramming, which is intimately controlled by histone modifications, and (2) loss of histone acetylation has been extensively documented in human cancer (66,67).

There are various mechanisms by which HDAC inhibitors promote anti-tumor activity, including cell cycle arrest(68–74), apoptosis(72,75), and DNA damage response(76). Indeed, HDAC inhibitors have been shown to promote differentiation in hematologic and solid tumors. Oncogenic proteins complex with, co-opt, and aberrantly recruit HDACs to repress genes that obstruct tumorigenesis(25,77), such as those regulating differentiation, which can be reversed by HDAC inhibitors(78,79). Blocking HDAC6 and HDAC3 in glioblastoma and glioma stem cells, respectively, promoted differentiation and reduced self-renewal in preclinical models (80,81). In neuroblastoma, HDAC8 inhibition alone or in combination with retinoic acid led to cell cycle arrest and differentiation in cancer cell lines and xenografts(82,83); similarly, HDAC1/2 inhibition, alone or with retinoic acid, also induced differentiation and decreased viability in neuroblastoma cell lines(84). HDAC inhibition promotes differentiation of AML via different molecular pathways(85,86). A unifying mechanism underlying differentiation induction by HDAC inhibition among cancers has yet to be proposed.

In our study, HDAC1/2 inhibition resulted in induction of broad differentiation signals that were associated with and partially dependent on H3K27ac. HDAC1/2 inhibitors exerted differential effect on various acetyl positions of histone proteins, consistent with prior reports(87,88). Despite extensive histone acetylation by HDAC1/2 inhibition, only H3K27ac at HDAC1/2-bound regions associated with a differentiation gene expression profile. Previous studies have linked different histone acetylation marks with CRC prognosis and histological subtypes(89–91). H3K27ac, a marker of active enhancers, had a distinct binding pattern when comparing CRC to native colon tissues(92,93). Importantly, degradation of EP300, a principal acetyltransferase of H3K27, partially reversed the differentiation phenotype induced by HDAC1/2 inhibition, ascribing functionality to this mark in promoting differentiation. This raises the possibility that a fine balance between EP300 and HDAC1/2 may govern stem cell and differentiation programs in normal intestinal homeostasis as well as CRC. Finding therapeutic options that selectively activate H3K27ac without impacting other histone PTMs is one potential approach to bypass HDAC1/2 inhibitor toxicity and sustain anti-tumor activity.

In addition to directly activating differentiation genes via gains in H3K27ac, HDAC1/2 inhibition also transcriptionally induced expression of DAPK3. As a functional participant in p53 signaling and therapy resistance, DAPK3’s role in tumor biology is gaining recognition(94–97). We found that genetic or pharmacological inhibition of DAPK3 partially reverses differentiation induction by HDAC1/2 inhibition. The phospho-proteome regulated by DAPK3 and its mechanistic role in CRC differentiation requires further elucidation.

Drug toxicity may limit the ability to achieve therapeutic levels of HDAC inhibitors required to induce tumor differentiation, potentially explaining the suboptimal clinical trial results. By combining selective HDAC1/2 inhibitors with chemotherapeutics that are active in CRC, like 5-FU, we may limit systemic levels, reduce toxicity, and enhance anti-tumor responses(98). Preclinical studies in solid tumor cell lines showed synergism or potentiation of HDAC inhibitors with various agents(99). In particular, Vorinostat was shown to synergize with 5-FU(100), oxaliplatin(101), and irinotecan(102). Existing targeted therapies have limited response in CRC compared to other cancers, likely due to the contribution of cancer cell plasticity. For example, CRC resistance in response to newly developed KRAS inhibitors is mediated by activation of EGFR signaling(2,4,7,8), which is an intrinsic transcriptional and/or epigenetic adaptive response. Since HDAC1/2 inhibition and epigenetic reprogramming can restrict plasticity by promoting differentiation, combining HDAC1/2 inhibitors with existing targeted therapies may improve anti-tumor responses by preventing or delaying resistance mediated by tumor plasticity. Indeed, a recent study showed that inhibition of histone methyltransferase EZH2 synergized with RAS pathway inhibitors to promote CRC differentiation and regression in preclinical models(103). Interestingly, combining HDAC inhibitor chidamide with VEGF monoclonal antibody bevacizumab sensitized patients with unresectable chemotherapy-refractory locally advanced or metastatic microsatellite stable CRC to PD-1 checkpoint blockade in a phase II trial (104). Another strategy to reduce systemic toxicity is to restrict circulating levels and enrich anti-tumor concentrations of HDAC inhibitors by conjugating to antibodies imbued with CRC specificity. Finally, identifying predictive biomarkers to identify patients with CRC that will selectively respond to HDAC1/2 inhibitors may also increase chances for clinical translation.

## STUDY LIMITATIONS

While we narrowed our search of HDAC1/2 inhibitor-mediated PTMs to histone proteins, non-histone proteins are also known to be modified by HDACs(105) and can be dysregulated in cancer(106); we therefore cannot rule out their contribution to the differentiation phenotype observed. It is difficult to divorce cell cycle regulation from differentiation programs. While we observe upregulation of differentiation markers prior to reduced proliferation phenotypes, we cannot definitively conclude how these related cellular programs interact during HDAC1/2 inhibition in CRC.

## METHODS

### Inactive MRK60 synthesis

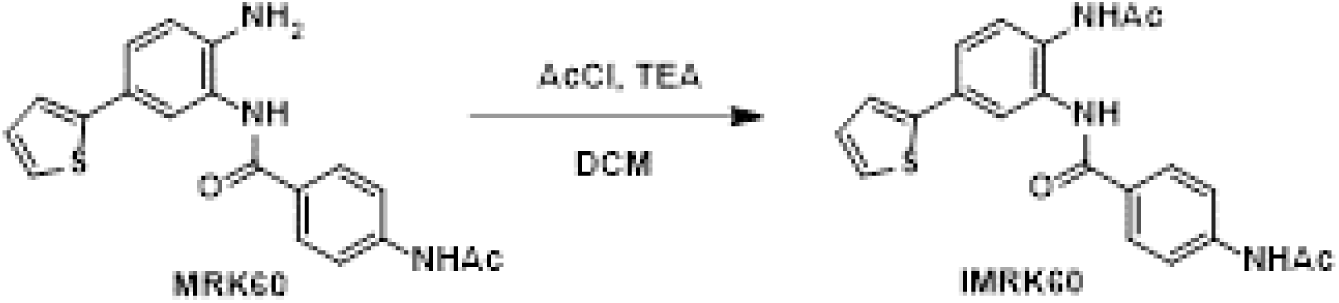

In a 4 mL vial fitted with a stir bar, **MRK60** (10 mg) was dissolved in dichloromethane (100 uL), followed by addition of triethylamine (TEA, 20 uL) and acetyl chloride (AcCl, 7 uL). The reaction mixture was stirred at room temperature overnight. The resulting mixture was concentrated in vacuo and purified by C18 reverse-phase column chromatography (eluent: water/acetonitrile, 0.1% formic acid) to give 6.55 mg of **inactive MRK60** (**iMRK60)** as white powder (Yield: 59 %).

^1^H NMR (500 MHz, DMSO-*d*_6_) δ 10.24 (s, 1H), 9.83 (s, 1H), 9.74 (s, 1H), 7.94 (s, 1 H), 7.93 (d, *J* = 8.6 Hz, 2H), 7.73 (d, *J* = 8.8 Hz, 2H), 7.58 (d, *J* = 8.4 Hz, 1H), 7.54 (dd, *J* = 5.0, 1.2 Hz, 1H), 7.50 (dd, *J* = 8.4, 2.2 Hz, 1H), 7.47 (dd, *J* = 3.6, 1.2 Hz, 1H), 7.14 (dd, *J* = 5.1, 3.6 Hz, 1H), 2.104 (s, 3H), 2.095 (s, 3H);

MS (ESI) calculated. For C_21_H_20_N_3_O_3_S [M+1]^+^ : 394.12, Found: 394.16.

### Synthesis of Biotin-MRK60

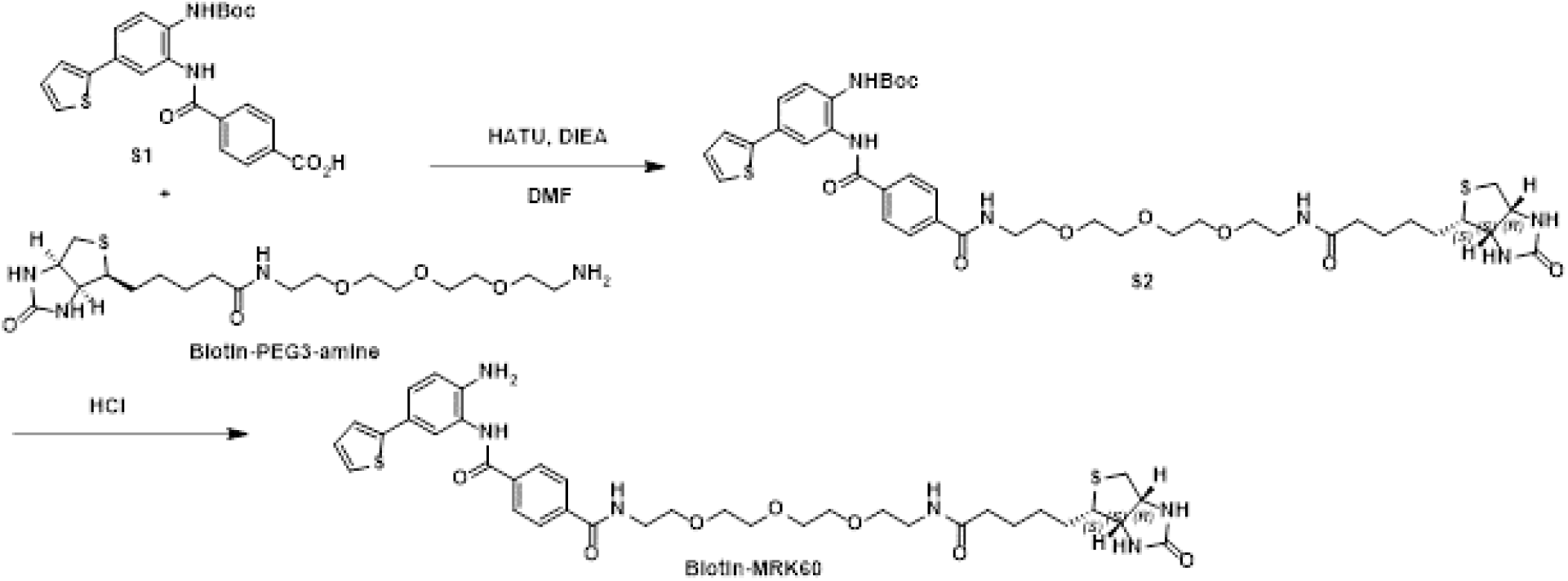

In a 4 mL vial fitted with a stir bar, **S1** (8.8 mg), **Biotin-PEG3-amine** (8.4 mg), and HATU (8.4 mg) were dissolved in 100 uL DMF. Then DIEA (18 uL) was added and stirred at room temperature for 2 h. The resulting mixture was concentrated under a steam of nitrogen and further purified by C18 reverse-phase column chromatography (eluent: water/acetonitrile, 0.1% formic acid) to give 17.37 mg of **S2** as white powder.

^1^H NMR (500 MHz, DMSO-*d*_6_) δ 10.01 (s, 1H), 8.77 (s, 1H), 8.71 (t, *J* = 5.6 Hz, 1H), 8.06 (d, *J* = 8.5 Hz, 2H), 8.00 (d, *J* = 8.5 Hz, 2H), 7.87 – 7.79 (m, 2H), 7.63 (d, *J* = 8.5 Hz, 1H), 7.55 – 7.50 (m, 2H), 7.46 (dd, *J* = 3.4, 1.2 Hz, 1H), 7.13 (dd, *J* = 5.0, 3.7 Hz, 1H), 6.40 (s, 1H), 6.35 (s, 1H), 4.29 (d, *J* = 5.0 Hz, 1H), 4.11 (ddd, *J* = 7.8, 4.4, 1.9 Hz, 1H), 3.60 – 3.43 (m, 13H), 3.17 (q, *J* = 5.8 Hz, 2H), 3.08 (ddd, *J* = 8.5, 6.1, 4.4 Hz, 1H), 2.80 (dd, *J* = 12.5, 5.1 Hz, 1H), 2.57 (d, *J* = 12.4 Hz, 1H), 2.54 (s, 1H), 2.06 (t, *J* = 7.4 Hz, 2H), 1.65 – 1.39 (m, 4H), 1.45 (s, 9 H), 1.35 – 1.21 (m, 2H). MS (ESI) calculated. For C_41_H_55_N_6_O_9_S_2_ [M+1]^+^ : 839.35, Found: 839.43.

In a 4 mL vial fitted with a stir bar, **S2** (17.37 mg) was dissolved in DCM (200 uL) followed by dropwise addition of HCl solution (4.0 M in dioxane, 100 uL). The reaction was stirred at room temperature for 1 h, and the solvent was removed. The residue was purified by C18 reverse-phase column chromatography (eluent: water/acetonitrile, 0.1% formic acid) to give 4.62 mg of **Biotin-MRK60** as white powder (Two steps yield = 31%).

^1^H NMR (500 MHz, DMSO-*d*_6_) δ 9.86 (s, 1H), 8.69 (t, *J* = 5.6 Hz, 1H), 8.07 (d, *J* = 8.2 Hz, 2H), 7.97 (d, *J* = 8.1 Hz, 2H), 7.83 (t, *J* = 5.7 Hz, 1H), 7.48 (d, *J* = 2.2 Hz, 1H), 7.35 (d, *J* = 5.1 Hz, 1H), 7.31 (dd, *J* = 8.3, 2.2 Hz, 1H), 7.25 (d, *J* = 3.5 Hz, 1H), 7.05 (dd, *J* = 5.1, 3.5 Hz, 1H), 6.82 (d, *J* = 8.4 Hz, 1H), 6.40 (s, 1H), 6.34 (s, 1H), 5.18 (s, 2H), 4.29 (dd, *J* = 7.8, 5.0 Hz, 1H), 4.11 (ddd, *J* = 7.4, 4.5, 1.9 Hz, 1H), 3.59 – 3.43 (m, 14H), 3.17 (q, *J* = 5.9 Hz, 2H), 3.08 (ddd, *J* = 8.7, 6.1, 4.3 Hz, 1H), 2.80 (dd, *J* = 12.4, 5.1 Hz, 1H), 2.57 (d, *J* = 12.4 Hz, 1H), 2.06 (t, *J* = 7.4 Hz, 2H), 1.65 – 1.38 (m, 4H), 1.35 – 1.19 (m, 2H);

MS (ESI) calculated. For C_36_H_47_N_6_O_7_S_2_ [M+1]^+^ : 739.29, Found: 739.21.

### Synthesis of Biotin-iMRK60

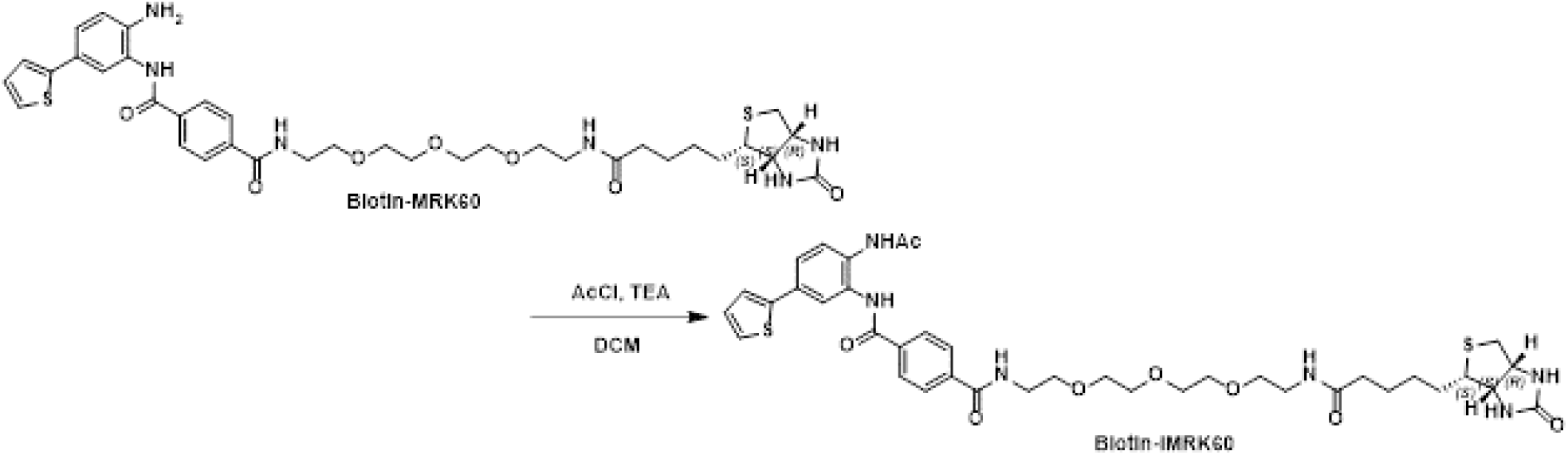

In a 4 mL vial fitted with a stir bar, **Biotin-MRK60** (4.62 mg) was dissolved in dichloromethane (5 uL), followed by addition of triethylamine (TEA, 5 uL) and acetyl chloride (AcCl, 3 uL). The reaction mixture was stirred at room temperature for 2 h. The resulting mixture was concentrated in vacuo and purified by C18 reverse-phase column chromatography (eluent: water/acetonitrile, 0.1% formic acid) to give 4.35 mg of **Biotin-iMRK60** as white powder (Yield: 59 %).

^1^H NMR (500 MHz, DMSO-*d*_6_) δ 10.06 (s, 1H), 9.75 (s, 1H), 8.70 (t, *J* = 5.6 Hz, 1H), 8.06 (d, *J* = 8.2 Hz, 2H), 7.99 (d, *J* = 8.4 Hz, 2H), 7.92 (d, *J* = 2.2 Hz, 1H), 7.82 (t, *J* = 5.7 Hz, 1H), 7.62 (d, *J* = 8.4 Hz, 1H), 7.57 – 7.50 (m, 2H), 7.48 (d, *J* = 3.4 Hz, 1H), 7.14 (dd, *J* = 5.1, 3.6 Hz, 1H), 6.40 (s, 1H), 6.34 (s, 1H), 4.29 (dd, *J* = 7.8, 5.1 Hz, 1H), 4.11 (ddd, *J* = 7.4, 4.4, 1.9 Hz, 1H), 3.60 – 3.43 (m, 14H), 3.17 (q, *J* = 5.9 Hz, 2H), 3.08 (ddd, *J* = 8.6, 6.1, 4.3 Hz, 1H), 2.80 (dd, *J* = 12.5, 5.1 Hz, 1H), 2.57 (d, *J* = 12.4 Hz, 1H), 2.09 (s, 3H), 2.05 (t, *J* = 7.3 Hz, 2H), 1.65 – 1.38 (m, 4H), 1.35 – 1.19 (m, 2H); MS (ESI) calculated. For C_38_H_49_N_6_O_8_S_2_ [M+1]^+^ : 781.30, Found: 781.52.

### Cell culture

All cell lines were maintained at 37 °C with 5% CO2. The human colorectal cancer cell lines HT115 (RRID:CVCL_2520), HT29 (HTB-38; RRID:CVCL_0320) and LS180 (CL-187; RRID:CVCL_0397) and HEK293T (RRID:CVCL_0063) were obtained from CCLE core facility at the Broad Institute, MIT and used at early passage for the experiments. Cells were maintained in DMEM medium supplemented with 10% FBS and 1% penicillin/streptomycin. Mycoplasma testing was performed every 3 months and found to be negative on each check.

### Endogenous reporter cell lines

Endogenous reporter cell lines were adopted from prior study (29). HT115^KRT20-mKate2^, and LS180^SOX9-GFP^, HT29^KRT20-GFP^, HT29^KRT20-GFP^ single reporters, and HT29^SOX9-mKate2/KRT20-GFP^ dual reporter were used in this study.

### Chemical screen targeting epigenetic regulators

We assembled a small library from selected members of epigenetic regulator families (Histone Acetylation (BRD), Histone deacetylation (HDAC & SIRT), Histone Lysine Methyltransferase (SET & KMT), Chromatin remodeling BAF(SWI/SNF) complex, and DNA Methyltransferases (DNMT)). All drugs were treated at 10 μM except for Pomalidomide at 20 μM for 48 hours. Cells were then stained with DAPI without fixation prior to the capture of fluorescent signal by the Celigo^TM^ High Throughput Micro-Well Image Cytometer. Please see Table S1 for complete list.

### Patient-derived Organoids

A tumor specimen from an unidentifiable patient with CRC was collected post colectomy under approval (protocol 13-189 and 14-408) by the Internal Review Board of the Dana Farber Cancer Institute, Boston, Massachusetts, USA. Notable genomic alterations in the patient-derived CRC organoid includes APC: c.835-8A>G (pathogenic intronic splice variant, rs1064793022) and KRAS: G12V.

Adenoma tissue from an unidentifiable patient with FAP were collected after colectomy under approval (protocol 13-189) by the Internal Review Board of the Dana-Farber Cancer Institute, Boston, MA, USA.

For generation of organoids from the patient-derived and mouse-derived tissues, the tissues were treated with EDTA and then resuspended in 30–50 μl of Matrigel (BD Bioscience) and plated in 24-well plates. WNT/R-spondin/Noggin (WRN) containing DMEM/F12 with HEPES (Sigma-Aldrich) containing 20% FBS, 1% penicillin/streptomycin and 50 ng/ml recombinant mouse EGF (Life Technologies) was used for culturing colon organoids. For the first 2–3 days after seeding, the media was also supplemented with 10 mM ROCK inhibitor Y-27632 (Sigma Aldrich) and 10 mM SB431542 (Sigma Aldrich), an inhibitor for the transforming growth factor (TGF)-β type I receptor to avoid anoikis. For passage, colon organoids were dispersed by trypsin-EDTA and transferred to fresh Matrigel. Passage was performed every 3-4 days with a 1:3–1:5 split ratio. For human colon organoid culture, the previous media was supplemented with antibiotics 100 μg/ml Primocin (Invivogen), 100 μg/ml Normocin (Invivogen); serum-free supplements 1× B27 (Thermo Fisher (Gibco)), 1X N2 (Thermo Fisher (Gibco)); chemical supplements 10 mM Nicotinamide (Sigma), 500mM N-acetylcysteine (Sigma), hormone 50 mM [Leu15]-Gastrin (Sigma), growth factor 100 μg/ml FGF10 (recombinant human) (Thermo Fisher) and 500nM A-83-01 (Sigma), which is an inhibitor of the TGF-β Receptors ALK4, 5, and 7.

### Ethical statement

Protocols were approved by the Internal Review Board of the Dana Farber Cancer Institute, Boston, Massachusetts, USA (protocols 13-189 and 14-408). Written consent was obtained from all participating patients, which included consent to publish results. All mice and experimental protocols were approved by Institutional Animal Care and Use Committee (IACUC) of Dana-Farber Cancer Institute (11-009). The maximal tumor size permitted by the IACUC is 2 cm, which was not exceeded.

### Animal studies

For in vivo studies, MRK60 was prepared with 10% DMSO, 45% PEG-400, and 45% sterile water to a final concentration of 6.25-12.5 mg/mL.

The generation of the *Lgr5^CreERT2^; Apc^f/f^; R26^tdT^* mouse was described earlier (14,28), mice aged 10-12 weeks were induced with tamoxifen at 2 mg/25 g on Day 0. MRK60 was injected intraperitoneally at 50 mg/kg dose with 2 days ON - 1 day OFF - 2 days ON - 2 days ON schedule from Day 10-24. When the mouse died or met the euthanasia requirement, bowels were harvested. Fresh tissue was collected for RNA isolation and protein collection. The rest of the bowels were fixed in formalin as above.

For *Lgr5^CreERT2^; Apc^f/f^; Kras^G12D^* mouse model, the *Lgr5^CreERT2^; Apc^f/f^; R26^tdT^* mice were bred with *Kras^LSL-G12D^* mice. Mice aged 10-12 weeks were induced with tamoxifen at 2 mg/25 g on Day 0. When the mouse died or met the euthanasia requirement, bowels were harvested and adenoma were dissociated into single cell suspension for FACS. Sorted DAPI^-^, tdTomato^+^ cells were established as organoids as below.

For tumor xenografts, 1 million *Apc^f/f^; Kras^G12D^* organoid cells were injected into flanks of athymic Ncr-nu/nu mice (RRID:IMSR_JAX:002019). MRK60 treatment was initiated 3 weeks after organoid injection to enable xenograft establishment. The treatment schedule was intraperitoneal injections once daily for 10 days at 50 mg/kg dose. Tumor volumes were obtained on every 3–4 days using handheld calipers to measure tumor length, width, and height to calculate cubic millimeters; mice were weighed using a digital balance. Tumor volumes were calculated using the following formula: volume = length × width^2^ × 0.5. Tumors were harvested, fixed in 10% formalin overnight, and paraffin-embedded for histologic analysis. Fresh tissue was also collected for RNA isolation, protein collection, and flash-frozen for long-term storage.

For the *Cdx2^CreERT2^; Apc^f/f^; R26^tdT^* mouse model, *Cdx2^CreERT2^* mice (RRID:IMSR_JAX:022390) were crossed with *Apc^f/f^; R26^tdT^* mice. Mice aged 10-12 weeks were induced with tamoxifen at 2 mg/25 g on Day 0. When the mouse died or met the euthanasia requirement, bowels were harvested and adenoma were dissociated into single cell suspension for FACS. Sorted DAPI^-^, tdTomato^+^ cells were established as organoids as below.

### CellTiter-Glo**®** cell viability assay

Cell and organoid viability were measured by CellTiter-Glo® Luminescent Cell Viability Assay and CellTiter-Glo® 3D Cell Viability Assay, respectively. Cells were plated in a flat-bottom white 96-well plate. Old media was removed to have 100 uL left. 50 uL of CTG solution was added and incubated for 10 minutes at room temperature on the rocker. Luminescent signal was measured by BMG LABTECH, FLUOstar Omega, SN 415-3943.

### ZIP synergy score

Viability was calculated by number of DAPI positive cells in each condition compared to the DMSO control in triplicate. ZIP synergy score was calculated with an R package “SynergyFinder” (RRID:SCR_019318) (42).

### Flow cytometry and cell sorting (FACS)

FACS was performed on the FACSAria III platform (BD) at DFCI Flow-core facility. For GFP, 488 nm laser and 530/30 filter were used. For mKate2, 561 nm laser and 610/20 filter were used. For tdTomato, 561 nm laser and 582/15 filter were used.

### Immunofluorescent imaging

Cells were seeded in the Corning flat-bottom dark 96-well plate in replicates. After drug treatment for 3 days, cells were fixed with 3% paraformaldehyde in PBS for 10 minutes at room temperature and washed with PBS 3 times. Cell membranes were permeabilized with 0.1% Triton X-100 in PBS 100 uL for 15 minutes at room temperature and washed with cold PBS 3 times. 2% BSA in PBST (PBS with 0.1% Tween 20) 100 uL was added as a blocking buffer and was incubated for 1 hour at room temperature. Primary antibodies were incubated at 1:500 dilution in a 4°C cold room rocker overnight and washed with cold PBST 3 times. Secondary antibodies were incubated at 1:2,000 dilution for 1 hour at room temperature and washed with cold PBST 3 times. DAPI was added at 1:2,000 in PBST for 10 minutes before washing with cold PBST. Images were captured by fluorescent microscopy using the Olympus Life Sciences IX73 microscope at 40x magnification. In separate experiments where varied doses of treatment were done with endogenous fluorescent reporter cells, fluorescent signal was captured by Celigo^TM^ High Throughput Micro-Well Image Cytometer.

### Histopathology

Paraffin-embedded intestines and xenograft tumors were serially sectioned and mounted on a microscopic glass slides. Sections were subjected to hematoxylin and eosin (H&E) as well as immunohistochemistry, using standard procedures. For morphological analysis, sections were serially dehydrated in xylene and ethanol and stained with H&E for histological assessment or KRT20. Antibody used is listed in Table S2.

### Immunoblot

Cells were lysed in RIPA buffer supplemented with a protease inhibitor cocktail (Roche) and phosphatase inhibitor (Cell Signaling 5870s). Protein were measured with Pierce^TM^ Rapid Gold BCA Protein Assay Kit (Thermo Fisher A53227) and denatured in 4× Laemmli buffer (Biorad 1610747). Whole cell extracts were resolved by 8–16% Tris-glycine polyacrylamide gel (Invitrogen XP08165BOX), transferred to PVDF membranes with iBlotTM 2 transfer device (Thermo Fisher IB21001) using P0 protocol, and probed with indicated primary antibodies. Bound antibodies were detected with LI-COR IRDye ® 680/800CW anti-rabbit/mouse secondary antibodies. Primary antibodies are listed in Table S2.

### RNA isolation, RT-qPCR, and RNA-sequencing

Total RNA was isolated using the RNeasy Mini Kit (Qiagen 74004) and cDNA was synthesized using the iScriptTM Reverse Transcription Supermix for RT-qPCR (Bio-Rad 1708840). Gene-specific primers for SYBR Green real-time PCR were synthesized by Integrated DNA Technologies. RT-qPCR was performed and analyzed using CFX96 Real-Time PCR Detection System (Bio-Rad 1845096) and using Power SYBR Green PCR Master Mix (Thermo Fisher 4368577). Relative mRNA expression was determined by normalizing to ACTB expression.

For RNA-sequencing, total RNA was isolated as above, before submission to Novogene Corporation Inc. for eukaryotic mRNA-seq on the NovaSeq6000 (paired-end 150-base-pair sequencing) with poly-A capture, non-directional library preparation. 10 Gb per sample was targeted.

### RNA-sequencing preprocessing and data analysis

Raw sequence quality was assessed using fastp v0.23.4(107). More than 35 million reads of each RNA-seq sample were aligned to human genome assembly GRCh38 using STAR v2.7.11(108). Only unique alignments to the genome were allowed. Read counts for each gene is generate by RSEM v1.3.3(109). The differential expression (DE) analyses were performed using DESeq2 v1.44.0(110). Reads for each sample were normalized by the DESeq2 method of median of ratios. The significance level was set at 0.05 for false discovery rate (FDR). Gene set enrichment analysis for DEGs was performed using Gene Set Enrichment Analysis (GSEA v4.3.3) (RRID:SCR_003199) (111).

### Histone extraction

Histone was extracted from cell lysates with EpiQuik™ Total Histone Extraction Kit. Histone protein then proceeded with the same protocol as the above immunoblot protocol. H3 was used as a housekeeping gene.

### Histone profiling

HT29 cells were grown and treated with DMSO, MRK60 and iMRK60 for either 24 or 48 hours. Cells were pelleted and histones were extracted following an established protocol(112). The purity of extracted histones was assessed using an SDS-PAGE. Briefly, samples (each containing 10 μg of histones) were propionylated, desalted and digested overnight with trypsin. A second round of propionylation occurred at the peptide level and peptides were desalted using C18 Sep-Pak cartridges (Waters). A reference mixture of isotopically labeled synthetic peptides for histones H3 and H4 was added to each sample prior to MS analysis. Peptides were separated on a C18 column (EASY-nLC 1000, Thermo Scientific) and analyzed on a Q Exactive^TM^ Plus (Thermo Scientific) using a PRM method. Peak areas were extracted in Skyline(113) and integrated. The ratios of endogenous to heavy peak areas were log2 transformed and normalized to unmodified regions (“Norm” peptide) of histone H3 or H4 respectively. Samples were normalized to DMSO controls (for each histone mark, the DMSO median was subtracted from each sample). Similarity matrices were generated using Spearman rank correlation across samples. Detailed protocols for sample preparation steps can be found at https://panoramaweb.org/labkey/wiki/LINCS/Overview%20Information/page.view?name=sops.

### Assay for transposase-accessible chromatin sequencing (ATAC-seq)

ATAC libraries were prepared as described previously(114). Briefly, 25,000 cells in duplicates were lysed to prepare nuclear pellets, which then underwent transposition with TDE1 Enzyme (Illumina, 20034197). Tagmented DNA was purified using Zymo DNA Clean and Concentrator-5 Kit (catalog no. D4014), and the purified DNA was PCR-amplified with NEBNext 2X Master Mix and Illumina adapters. The libraries were purified after PCR using AMPure XP beads (Beckman Coulter). 150-bp paired-end reads were sequenced on a NovaSeq instrument (Illumina).

### ATAC-seq data processing

ATAC-seq data were analyzed as described previously(115). Briefly, reads were mapped to the GRCh38 human genome using Bowtie2 aligner (version 2.3.5) (RRID:SCR_016368) (116). Model-based Analysis of ChIP-Seq (MACS2) (RRID:SCR_013291) was used to call peaks using the parameters “-t $bamfile -f BAMPE -n qc/macs/$base.macs2 -q 0.01 -g mm”(44,117). Bigwig files were generated using deepTools (RRID:SCR_016366) bamCoverage function with the options “bamCoverage --binSize 10 --smoothLength 30 -p 4 --normalize using RPGC --effectiveGenomeSize 2730871774 --extendReads $fragLength -b $file -of bigwig” (118). All bigwig and bed files were filtered using the ENCODE Blacklist. Only peaks with < 0.00001 were considered for further analyses. Proximal peaks (−2 kb to 2 kb relative to transcription start site (TSS) called by MACS2 were linked/annotated to genes by ChIPseeker (version 1.34.1) (RRID:SCR_021322) (119). Then, DiffBind (version 3.14.0) (RRID:SCR_012918) (120) were used to visualize peaks.

### Cleavage Under Targets and Release Using Nuclease sequencing (CUT&RUN)

CUT&RUN was performed in accordance with the Epicypher® CUT&RUN kit (Epicypher, #14-1048) and the kit manual version 3.1 with the following specifications. Buffers were prepared with Triton X-100 1% and SDS 0.05% and digitonin 0.01%. Cells were treated by either DMSO, MRK60 2 µM, iMRK60 2 µM, and Romidepsin 0.005 nM for 48 hours. The media was removed and fixing solution (DMEM and PFA 1%) 900 uL was added for 1 minutes before quenching with 1.25 M glycine 100 uL to a final concentration of 125 mM. Cells were collected with cell scraper and were centrifuged at 600 g for 3 minutes at room temperature. Pellets were collected and redistributed in cold PBS at a final concentration of 500,000 cells/ sample and proceeded with the protocol step 7. There were 2 technical replicates per condition. SNAP-CUTANA Spike-in control 2 µL were added in the IgG control tubes. The antibodies used for binding are listed in Table S2. There were incubated overnight in a 4°C room on a nutator. pAG-MNase and target chromatin digestion and release per protocol. E. coli spike-in DNA 1 µL was added at the end. Reverse crosslink was done by adding 10% SDS and proteinase K and incubated at 55°C for 12 hours in a thermocycler. The DNA was then isolated and diluted and then quantified using Qubit dsDNA HS assay kit (Agilent Technologies). Library preparation was done with NEBNext Ultra II DNA prep kit and 96 unique dual index primer pairs (NEBNext® E6440S kit) with a few modifications. Briefly, end repair and dA-tailing were conducted on 5 ng of CUT&RUN eluted DNA for 30 minutes at 20°C followed by 30 minutes hour at 65°C. After adaptor ligation for 30 min at 20°C, the DNA fragments were purified by 0.9× volume of AMPure XP beads (Beckman Coulter) followed by 14 cycles of PCR amplification with Next Ultra II Q5 master mix. The PCR products were purified with 1× volume of AMPure XP beads. After quantitative and qualitative analysis, libraries with different indexes were pooled and sequenced on Illumina NovaSeq platform with paired-end 150-bp reads.

### CUT&RUN data processing

CUT&RUN sequencing data were analyzed as described previously(121), following a standard pipeline (https://yezhengstat.github.io/CUTTag_tutorial/index.html). Briefly, paired-end 150-bp reads were aligned to GRCh38 human genome using Bowtie2 version 2.2.5(116) with the following options: --local --very-sensitive --no-mixed --no-discordant --phred33 -I 10 -X 700. Technical replicates (n = 4 per condition) were merged before read alignment to increase the power of peak calling. Model-based Analysis of ChIP-Seq (MACS2) (117) was used to call peaks from bam files. For SOX9 CUT&RUN peak calling, parameters —t input_file –p 1e-5 –f BAM –keep-dup all –n out_name was used to call narrow peaks. To check the SOX9 binding profile and enhancer activity of specific gene sets, proximal peaks (−2 kb to 2 kb relative to TSS) called by MACS2 were linked/annotated to genes by ChIPseeker version 1.34.1(119). Then, DiffBind version 3.14.0(120) were used to visualize peaks.

### Chem-sequencing (Chem-seq)

Chem-seq is a ChIP-based method to identify the sites bound by small chemical molecules throughout the human genome. The protocol is adapted from Anders, et al. 2014(34). Biotinylated MRK60 and iMRK60 were synthesized for this experiment. Anti-biotin antibody was used to pull down the biotinylated compounds and their complex.

For the in-vivo Chem-seq, either biotinylated MRK60 or biotinylated iMRK60 10 uM were added for 1 or 2 hours in incubator before fixing the cells. For the in-vitro Chem-seq, the chromatin lysate after the sonication step was added with either biotinylated MRK60 or biotinylated iMRK60 10 uM and was incubated on a 4°C rocker overnight.

Cells were fixed with formaldehyde at final concentration of 1% for 10 minutes on a rocker at room temperature before quenching with 1.25 M Glycine at 10% of the media volume for 5 minutes. Cells were collected with cell scraper and were centrifuged at 600 g for 4 minutes at 4°C. Pellets were washed with cold PBS for 3 times. 1.1 mL of the sonication buffer was added and transfer to shearing tubes. Chromatin was sheared with a Covaris E220 machine with the following settings: Peak Incident Power 150, Duty Cycles 5%, Cycles per Burst 200, Time 300 seconds. Lysates were centrifuged at 13,000 g at 4°C for 5 minutes and the supernatants were collected. Pierce™ High Capacity Streptavidin agarose 75 uL per reaction was washed with PBS at room temperature 3 times before using. The agarose was mixed with the chromatin lysate 450 uL and incubate at room temperature on a rocker for 15 minutes. Washing, eluting DNA-protein complex, reverse crosslinking, and DNA isolation were done as previously described.

### ChIP-seq and Chem-seq Analysis

ChIP-seq analysis. The ChiLin pipeline 2.0.0(122) was used for quality control and pre-processing of the data. Burrows-Wheeler Aligner (version 0.7.17-r1188) (RRID:SCR_010910) (123) was used to map reads and Model-based Analysis of ChIP-Seq (MACS2) (v2.1.0.20140616) (117) for peak calling using default parameters. Briefly, to find the nearest gene for ChIP-seq peaks, we used ‘bedtools closest’ (124) to get the closest (it can be overlapping or nonoverlapping) gene between peak file and reference gene file (+/- 1kbTSS). We used MACS2 (v2.1.2) to call narrow peaks using default parameters, cut offs: fdr="0.01", keepdup="1", extsize="146". Cis-regulatory element annotation system (CEAS) (RRID:SCR_010946) (125) analysis is used to annotate resulting peaks with genome features. Differential analysis of peaks was determined by DESeq2 (RRID:SCR_015687) (110). Super-enhancers were called by ROSE (RRID:SCR_017390) (126) in H3K27ac ChIP-seq data. Genomic Regions Enrichment of Annotations Tool (GREAT) (RRID:SCR_005807) (127) was used to annotate peaks with their biological functions. Cistrome toolkit(128,129) was used to probe which factors might regulate the user-defined genes. Conservation plots were obtained with the Conservation Plot (version 1.0.0) tool available in Cistrome toolkit (RRID:SCR_000242). ChIP-seq data visualization. Normalized profiles corresponding to read coverage per 1 million reads were used for heatmaps and for visualization using the integrative genomics viewer (IGV) (RRID:SCR_011793) (130). Wiggle tracks were visualized using IGV. Heat maps were prepared using DiffBind (version 3.14.0)(120).

### Oligo design and cloning

The pCC_01 - hU6-BsmBI-sgRNA(E + F)-barcode-EFS-Cas9-NLS-2A-Puro-WPRE plasmid (Addgene #139086; RRID:Addgene_139086) was used for CRISPR knockout. Briefly the plasmid was digested with Esp3I (New England Biolabs, R0734) at 37 °C overnight. The 13 kb band was purified with gel purification kit (Takara Nucleospin® Gel and PCR Clean-up Midi). The insert encoding the gRNA sequences was prepared by annealing TOP and BOTTOM oligonucleotides followed by T4 PNK phosphorylation (New England Biolabs, M0201S) in accordance with the following template: TOP: caccgNNNNNNNNNNNNNNNNNNNN, BOTTOM: aaacNNNNNNNNNNNNNNNNNNNNc. The purified backbone and insert were ligated together using T4 DNA Ligase (New England Biolabs, M0202S) following manufacturor’s protocols. **CRISPR sgRNA sequences**: HDAC1: ACTCCGACATGTTATCTGGA, HDAC2: GTATAGATGATGAGTCATAT, GCN1_g1: CCTTGTCGCTTGGCTCTCGA, GCN1_g2: GCAGTTCTGCACGAGTCACA.

### Lentivirus transfection

To generate lentiviruses, HEK293T cells were co-transfected with sgRNA expression vectors and lentiviral packaging constructs psPAX2 (Addgene #12260; RRID:Addgene_12260) and pMD2.G (Addgene #12259; RRID:Addgene_12259) in a 2:1:1 ratio using X-tremeGENE HP DNA Transfection Reagent (Roche) according to the manufacturer’s instructions. Cell culture media was changed the following day and lentiviral supernatant was harvested 24 h and 48 h later and filtered through a 0.45 μm filter (Celltreat). Lentiviruses were aliquoted and stored at −80°C until use.

To perform lentiviral infection, the CRC cells were plated in a 6-well flat bottom plate and infected with 0.5 mL virus in media containing 8 mg/mL polybrene overnight. The media was changed the next day. After 2 days, puromycin were started at 3 μg/mL respectively and continued until the negative control cells died.

### scRNA-seq in patient-derived CRC organoids

The patient-derived CRC organoid was treated with either MRK60 at 30 µM, JQAD1 20 µM, or both for 48 hours before the collection. At the time of collection, 5 mL of Cell Recovery Solution (Corning, 354253) was added for every well in a 6-well plate format and left in a rocker at 4°C for 20 minutes to dissolve the Matrigel. Organoids were collected in the 50 mL Falcon tube and spun down in a 4* centrifuge at 900g for 5 minutes. The supernatant was removed. The pellets (organoids) were mixed with 4 mL of prewarmed TrypLE Express (Thermo Fischer (Gibco)) Resuspended the pellet by pipetting up and down multiple times and incubates in a 37°C bead bath for 15 minutes with pipetting up and down every 5 minutes. Examined the samples under a microscope to ensure no clumping. Counted the cell using the Countess 3 microscope (Thermo Fisher) and diluted with 0.04% BSA in PBS to a concentration of 1,000 cells per µL. The organoids were then submitted to the single cell core at the Translational Immunogenomics Lab at Dana-Farber Cancer Institute, which prepared the library with a Chromium NEXT GEM Single Cell 3’ (v3.1 chemistry) kit. Each sample with 10,000 cells was sequenced by 150-base-pair, paired-end Illumina NovaseqX Plus sequencing with a target at 110Gb per 10,000 cells.

### scRNA-seq preprocessing, alignment, and gene counts

Demultiplexing, alignment to the transcriptome, and unique molecular identifier collapsing were performed using the Cell Ranger toolkit (version 7.1.0) (RRID:SCR_017344) provided by 10x Genomics. Standard procedures for quality control filtering, data scaling and normalization, detection of highly variable genes were followed using the Seurat (version 4) (RRID:SCR_016341) (131) in RStudio. Cells with unique feature counts lower than 100 and greater than 25,000 and cells with greater than 25% mitochondrial DNA were excluded. Counts were log-normalized and scaled by a factor of 10,000 according to the default parameters when using the Seurat LogNormalize function. Variable features were identified, and the data were scaled using the default parameters (Ngenes = 2000) of the FindVariableFeatures FIG and ScaleData Seurat functions, respectively. Principle components analysis (PCA) was completed on the remaining cells, and 10 principal components were selected for clustering, t-distributed stochastic neighbor embedding, and UMAP analyses. Cells were visualized primarily using UMAP nonlinear dimensional reduction (dimensions, 1:10; resolution = 0.3), from which feature plots were generated to demonstrate distribution of gene expression and MRK60 versus DMSO treated cells and expression levels of various marker genes throughout the population. Marker genes for each resulting cluster were found using the FindMarkers function with the minimum prevalence set to 25%. We implemented several quality control steps during data processing, including the use of CellBender(132) to remove ambient RNA contamination, DoubletFinder(133) to eliminate cell doublets, and Seurat(131)/Scanpy (RRID:SCR_018139) (134) data integration to correct for batch effect.

### Dataset integration

scRNA-seq IntegrateData function in Seurat v4(131) was used to counteract batch effects among different treatments. PCA was then completed on the integrated object, and the number of principal components selected for clustering was determined using the integrated object’s elbow plot. Cells were then visualized primarily using UMAP nonlinear dimensional reduction from which feature and violin plots were generated to demonstrate the distribution of gene expression and expression levels of various marker genes and gene signatures throughout the population.

### scRNA-seq gene signature analysis

To analyze existing gene signatures on our scRNA-seq data, the Seurat AddModuleScore function in Seurat v4(131) was used to calculate the average normalized and scaled gene expression of a given gene list in each individual cell. Specific cell types were identified using established marker genes and gene signatures (60). Gene signature scoring was then visualized with feature and violin plots. To generate novel gene signatures, the Seurat FindMarkers function was used to create lists of genes differentially expressed in one specified subset in comparison to another given subset. The minimum prevalence was set to 25%.

### Drug response trajectory analysis

After Scanpy(134) processing, single-cell transcriptomic profiles of the cells belonging to the colonocyte cluster were fitted using the diffmap() function. neighbors() was applied to build a graph based on diffusion map result (n_neighbors=10). By assuming that the similar cells would most resemble each other. Then all cells were visualized using draw_graph() function.

### Statistical analysis and reproducibility

Experiments were performed in triplicate. Data are represented as means ± SD unless indicated otherwise. For each experiment, either independent biological or technical replicates are as noted in the figure legends and were repeated with similar results. Statistical analysis was performed using Microsoft Office, Prism 9.0 (GraphPad) (RRID:SCR_002798), or RStudio (RRID:SCR_000432) statistical tools. Pairwise comparisons between groups (that is, experimental versus control) were performed using an unpaired two-tailed Student’s t test or Kruskal-Wallis test as appropriate unless otherwise indicated. For all experiments, the variance between comparison groups was found to be equivalent. Tests marked by “****”, “***”, “**”, and “*” represent they are statistically different from the controls of P values <0.0001, <0.001, <0.01, and <0.05, respectively. ns not significantly different. In all the boxplots, each box represents the interquartile range (IQR, the range between the 25th and 75th percentile) with the mid-point of the data, whiskers indicate the upper and lower value within 1.5 times the IQR.

### Data availability

All data that support the findings of this study are available within the paper and its supplemental files. Raw and processed sequencing data are available in the Sequence Read Archive (SRA) at the NCBI Center with the accession number PRJNA1142083 and Gene Expression Omnibus (GEO) with the accession number GSE273626. All processed data are also available at https://doi.org/10.5281/zenodo.13145655. Source data are provided with this paper. All other data supporting the findings of this study are available from the corresponding author on reasonable request.

### Code availability

All bioinformatic analysis tools and pipelines used in this study are documented in the method section. Codes are available from the corresponding author upon reasonable request.

### Author contributions

P.L., Z.L., and P.D. co-led the study. P.L. and P.D. performed most experiments, analyzed data, and wrote/revised the manuscript. Z.L. performed all bioinformatic analyses and wrote/revised the manuscript. S.S. performed chemical screen. A.K.R. performed validation and scRNA-seq experiment. Q.L., P.L., and S.L. curated chemical library, synthesized MRK60, inactive forms, and biotinylated forms, and synthesized EP300 degrader. D.C. provided bioinformatic support. P.B. and D.A. performed in vivo experiments. P.S. performed validation experiments. M.P., W.H., and S.A.C performed PTM histone profiling. H.P. provided clinical insight and revised manuscript. J.M.C provided clinical insight, provided funding, and revised manuscript. J.Q. co-supervised the study and revised manuscript. N.S.S. designed and co-supervised the study and wrote/revised the manuscript.

## Acknowledgements

We thank Sethi lab members, Rameen Beroukhim, and others for insightful discussions and/or review of the manuscript; Center for Cancer Epigenetics, Kornelia Polyak, Myles Brown, Ramesh Shivdasani, and Matthew Freedman for use of Celigo^TM^ High Throughput Micro-Well Image Cytometer; Christine Perret for kindly sharing the Apcflox/flox mice developed in her laboratory; Aniket Gad and Lay-Hong Ang for assistance with immunohistochemistry assays; Shuqiang Li and Kenneth Livak for assistance with scRNA-seq; Dana-Farber/Harvard Cancer Center for the use of the Specialized Histopathology Core, which provided histology and immunohistochemistry service; Harvard Digestive Disease Center and NIH grant P30DK034854 for core services, resources, technology, and expertise. Dana-Farber/Harvard Cancer Center is supported in part by an NCI Cancer Center Support Grant # NIH 5 P30 CA06516. This work was funded by the Colorectal Cancer Alliance, Virtual Scholar Award from the Department of Defense (CA201084), Bridge Project (DF/HCC-MIT), and 1R01CA292507 to N.S.S and philanthropic support from the Jimmy Fund Walk (Opiela Family), Craig Baskin, Dave & Carol Fischer, and Howard & Wendy Cox to the Sethi Lab.

Supplemental Information is available for this paper

Correspondence and requests for materials should be addressed to J.Q. and N.S.S.

